# Control of grain dimensions via ALLENE OXIDE CYCLASE

**DOI:** 10.64898/2026.01.15.699649

**Authors:** Jian Luo, Linsan Liu, Xiujuan Yang, Emily Lyon, Jonathan Griffin, Jennifer Shoesmith, Wenhao Wu, Ruth Hamilton, Mirjam Nuter, Mikah Katay-Fodor, Matthew Tucker, Sarah M McKim

## Abstract

Cereal grain is a global food staple. Grain formation starts with fertilization of the embryo sac which generates filial tissues, such as the embryo and endosperm, that grow embedded in the maternal ovule and surrounding ovary wall. Maternal tissues provide nutrition and protection but are progressively reduced and degraded, such that filial tissues dominate the mature grain. While we know some enzymatic players, regulation of this process in temperate cereals like wheat and barley, remains less understood. However, previous work suggested a role for the phytohormone Jasmonate (JA). To learn more, we edited the barley *ALLENE OXIDASE CYCLASE* (*HvAOC*) gene encoding a JA biosynthetic enzyme. Defective *HvAOC* alleles caused enhanced vegetative growth, faster flowering and larger, heavier grain - phenotypes rescued by MeJA treatment. HvAOC control of final grain parameters is parentally-derived, consistent with enriched *HvAOC* expression in maternal compared to filial tissues. Defective HvAOC function enlarged maternal tissues involved in nutrient transfer, correlating with larger endosperm and higher expression of genes linked to sucrose transport. This work provides the first genetic dissection of JA in barley and highlights a key role for HvAOC in the coordination between growth and death of maternal tissues with grain size and weight.

## Introduction

Grain from cultivated cereals provides more calories to the human diet than any other source. A complex multigenerational unit, grain originates from double fertilisation of the ovule’s embryo sac which generates the triploid endosperm and diploid embryo. These filial tissues grow embedded within the ovule’s nucellus and further enclosed by inner and outer integuments, which collectively make up the seed. In cereals, the seed is surrounded by the ovary wall or pericarp, forming the ‘caryopsis’ or grain fruit. While the embryo grows and differentiates, endosperm nuclei divide into a coenocyte which cellularises and fills with starch, driving caryopsis expansion between the lemma and palea hull (Olsen, 2020). At the same time, maternal tissues form nutritive and/or protective structures which ultimately die. For instance, the outer integuments disintegrate and the inner integuments and nucellar epidermis compress into a seed coat while pericarp cells proliferate, expand and subsequently crush into a protective case during grain filling (Kellogg, 2016; Brinton & Uauy, 2019). Nucellar tissues specialize into structures which facilitate nutrient transport into the filial tissues. For instance, wheat and barley, dominant temperate cereal crops, form the nucellar projection (NP) along the grain ventral axis which acts as a central conduit for assimilate and which later undergoes PCD close to the filial boundary (Theil *et al*., 2008; Hands *et al*., 2012; Dominguez & Cejudo, 2014). Collectively, these events shift the balance of tissues from the parental to the filial generation so that mature grain consists mainly of filial tissues held within reduced maternal tissues. Many alleles linked to changes in grain dimensions and/or weight also regulate ovary, nucellus, integument, pericarp and/or hull development (Yin & Xue, 2012; Ren *et al*., 2018; Zhao *et al*., 2018; Brinton & Uauy, 2019; Hao *et al*., 2021; Jia *et al*., 2021; Shoesmith *et al*., 2021; Yang *et al*., 2023; Long *et al*., 2024; Wang *et al*., 2024). However, the mechanistic links between genes regulating parental tissues and mature grain parameters are poorly understood. For instance, most agronomic alleles in rice reported to control grain size are only described to have a role in cell proliferation and/or expansion of hull organs (Riemann *et al*., 2013; Ren *et al*., 2018; Wang *et al*., 2024).

We recently demonstrated that the miRNA-regulated transcription factor HvAPETALA2 (HvAP2) controls specific events in parental differentiation and degradation in the grain of the widely cultivated temperate cereal barley (*Hordeum vulgare ssp. Vulgare*; Shoesmith *et al*., 2021). First characterized as a repressor of internode growth and developmental progression in the spike (Houston *et al*., 2013) and stem (Patil *et al*., 2019), HvAP2 also restricts grain and lemma length, limits integument layer number and accelerates maternal tissue degradation, especially of the NP, an activity linked to the expression of the MADS-Box gene *HvMADS29*, a key regulator of NP differentiation in barley and rice (Yin & Xue, 2012; Shoesmith *et al*., 2021). HvAP2-limitation of vegetative and grain development correlated with elevated jasmonate (JA)-responsive gene expression and sensitivity (Patil *et al*., 2019; Shoesmith *et al*., 2021), consistent with JA-related growth repression reported in many plants (Huang *et al*., 2017).

JA controls other developmental processes in plants, such as germination rate and flowering time, and plays a key signalling role in responding to biotic invaders and abiotic stresses (Wasternack & Hause, 2013; Campos *et al*., 2014; Machado *et al*., 2016; Huang *et al*., 2017; Li *et al*., 2025). Many insights about JA come from studying mutants defective in JA biosynthesis or signalling pathways which appear broadly conserved across land plants (Li *et al*., 2021). JA and its metabolites, collectively called jasmonates, are oxylipin phytohormones originating from plastid-derived linolenic acid which is first oxygenated by 13-lipoxygenase (13-LOX), then converted into an epoxide by a 13-allene oxide synthase (AOS), and further cyclized by an allene oxide cyclase (AOC) to cyclopentenone (cis)-12-oxophytodienoic acid (OPDA). Reduction by OPDA reductase (OPR) and three cycles of β-oxidation in peroxisomes produces (+)-7-iso-jasmonic acid (Wasternack *et al*., 2013) with further modifications yielding Jasmonoyl-isoleucine (JA-Ile), a major bioactive JA, as well as other active, inactive, or partially active compounds (Li *et al*., 2021; Zhu *et al*., 2021). Upon perception of bioactive JA, the coronatine insensitive 1 (COI1) receptor targets Jasmonate ZIM-domain (JAZ) proteins for degradation, releasing JAZ-mediated repression of selected transcription factors, especially the MYC2 transcriptional activator central to many JA responses (Wasternack & Kombrink, 2010; Wasternack & Strnad, 2018). Recent genetic dissection in Arabidopsis showed that JAZ6 promotes seed size while downstream activators in JA signalling repress integument cell proliferation and ovule size, roles associated reducing seed size (Hu *et al*., 2021). JA restriction of seed size may be conserved in monocots as both *coi* mutants and overexpression of *OsJAZ11* in rice increase grain width and weight (Lee *et al*., 2015; Mehra *et al*., 2022). However, our understanding about how JA controls reproductive and grain development remains fragmentary in cereals, especially in wheat and barley.

To address this, we gene edited the *AOC1* gene in barley. Predicted loss of HvAOC function alleles caused enhanced vegetative growth, accelerated flowering and enlarged, heavier grain. Grain dimension changes correlated with larger and more persistent maternal tissues involved in nutrient transfer as well as increased endosperm, but no change in hull organ size. These results reveal a key role for JA in post-fertilisation maternal tissue development and suggest that modulating JA levels may impact grain size and weight.

## Methods

### Germplasm and growth conditions

Barley plants for genotyping and phenotyping were grown in glasshouses (James Hutton Institute, UK) under long day photoperiod conditions (16 hr light 18°C/8 hr dark 14°C) in standard cereal compost mix: 1.2m^3^ Peat, 100l sand, 1.5kg Osmocote Exact Start, 3.5kg Osmocote Exact 3-4 month, 2.5kg Lime, Ca and Mg (each), 0.5kg Celcote, 100l Perlite and 280g Intercept. For germination, surface-sterilized seeds were germinated on moist filter paper in a controlled environment cabinet in 16h:8h light:dark cycle, set to provide a day photon irradiance of 300 μmol.m^−2^.s^−1^, 70% relative humidity, a day temperature of 18 °C and a night temperature of 18 °C. Germplasm details provided in Table S1.

### Guide design, transformation, selection and screening

Gene edited lines were generated in the Golden Promise cultivar by the John Innes Crop Transformation team. Two guides (Table S2) aligning to the 5’ end of the first exon of the *HvAOC* gene model (HORVU.MOREX.r3.6HG0619890) were cloned into the dual guide acceptor vector (L2\Hv\35-36) and transformed into Golden Promise as described in (Lawrenson & Harwood, 2018). Five T0 plants were screened and edits detected in one line (line 01-01). Other T0 lines did not show edits in this generation. All T0 plants were grown, seed harvested and replanted for T1 generation. T2 seed from eight T1 plants (01-01, 01-02, 02-01, 03-01, 04-01, 04-03, 11-01 and 12-01) were sown and screened for the presence of Cas9 and edits in *HvAOC*. Two T3 lines with predicted loss of function edits in *HvAOC* (n4 and n7) which were also Cas9-free were grown, confirmed by genotyping and selected for fine phenotyping. The presence of Cas9 and edited *HvAOC* were detected using primers targeted on CaMV 35S promoter and Cas9 protein region.

### Genotyping

Genomic DNA (gDNA) was extracted from young leaf tissue using the DNeasy Plant Mini Kit (Qiagen). Transgenic plants were genotyped for mutations in HvAOC and for the presence of the bcoCas9 transgene by PCR. HvAOC target sites were amplified using OneTaq 2× Master Mix with GC Buffer (NEB). Each 15 μl reaction contained 7.5 μl of 2× master mix, 1 μl of GC enhancer (NEB), 0.5 μl of each primer, 1 μl of gDNA template, and 4.5 μl of nuclease-free water. PCR cycling conditions consisted of an initial denaturation at 94°C for 5 min, followed by 35 cycles of 94°C for 30 s, 58°C for 30 s, and 68°C for 30 s. A final extension was performed at 68°C for 5 min. PCR products were purified with ExoSAP-IT (Applied Biosystems) prior to Sanger sequencing. Two primer pairs were designed to detect the presence of the *Cas9* gene. Each 10 μl PCR reaction contained 5 μl OneTaq 2× Master Mix with Standard Buffer (NEB), 0.2 μl of each primer, 1 μl gDNA, and 2.6 μl nuclease-free water. The cycling conditions consisted of an initial denaturation at 94°C for 30 s, followed by 35 cycles of 94°C for 30 s, 56°C for 30 s, and 68°C for 54 s, with a final extension at 68°C for 5 min. The *Zeo1.b* allele was screened using GoTaq® Flexi DNA Polymerase following the manufacturer’s recommendations. Each reaction contained 3 μl of 5× Green Flexi Buffer, 1.125 μl of MgCl₂, 0.75 μl of dNTP mix, 0.75 μl of each primer, 0.18 μl of GoTaq G2 enzyme, and 1 μl of gDNA template. PCR amplicons were digested with SfaNI (NEB) to identify the *Zeo1.b* allele. Primer sequences for all genotyping and *Cas9* transgene detection are listed in Table S3.

### Phenotyping

Coleoptile length was measured at 5 days after sowing (DAS) and germination rate recorded at 8 DAS. The number of tillers was counted at 32 DAS and 42 DAS. The DAS for anthesis was recorded for first three tillers. Anthesis (0 days after pollination, DAP) was defined as the stage when anthers turned yellow and released pollen upon gentle pressure with tweezers. Grains were sampled at 0, 6, 10, 12, 16 and 18 DAP, and grains from the central spike portion were photographed, weighed, and measured for length and width using ImageJ. Plant height, spike length, grain number for single spike and spike yield were measured at mature stage. Grain setting rate was calculated as the ratio of grain number to hull number. Grain dimensions and lemma length were determined with MARVIN-Universal (MARViNTECH GmbH, Germany) and digital calipers.

### Exogenous MeJA treatment

Exogenous MeJA (Sigma–Aldrich, USA, J2500) treatments were applied to both seedlings and flowering plants. A 10 mM MeJA stock solution was prepared with few drops of ethanol and diluted with water to the desired concentrations. For seedlings, MeJA or water (mock control) was sprayed from the stage of coleoptile emergence. Two concentrations of MeJA (0.1 and 0.25 mM) were applied every three days, for a total of four applications. Due to the small size of the seedlings, a portion of the spray solution contacted the soil surface. Coleoptile length was measured at 14 DAS. For flowering-stage plants, we followed previous studies in wheat and rice, and our earlier experiments in barley (Liu *et al*., 2012; Liu *et al*., 2017; Li *et al*., 2018; Patil *et al*., 2019), MeJA was applied at a concentration of 1 mM. Spraying was initiated at anthesis (0 DAP) and was repeated every three days until 12 DAP. During application, the pots were wrapped with plastic film to prevent spray solution from entering the soil. MeJA solution was evenly applied to the aerial parts of the plants, including leaves, stems, and developing spikes. Grains were sampled at 6 DAP, 18 DAP and at maturity. All spray solutions contained 0.01% (v/v) Tween-20 as a surfactant and water solution also contain same volume ethanol.

### Grain histology

Grains from Golden Promise and mutant lines were collected at 6, 10, 12, 18 DAP and fixed overnight in FAA, dehydrated in a series of 70, 80, 90 and 100% (v/v) ethanol and embedded in paraffin wax using Leica TP1020 Tissue Processor. Samples were sectioned to 10 μm thickness on microtome (Leica, RM2125). Sections were stained in 0.5% Toluidine blue (w/v) after dewaxing and imaged by digital slide scanner (ZEISS, Axioscan). For pericarp cell measurements, the ventral middle pericarp of fixed grain was dissected away manually, stained with 0.5% Toluidine blue and imaged with light microscope (Leica, MZ FLIII).

### Phylogenetic analysis

The barley *HvAOC* gene (HORVU.MOREX.r3.6HG0619890) was used as a query against the Ensembl Plants database (https://plants.ensembl.org/index.html) with default parameters to retrieve its homologues from other crop species, including monocots and eudicots. Phylogenetic relationships were inferred from amino acid sequences using a neighbour-joining method with 1,000 bootstrap replications. Conserved motifs among the proteins were explored using the MEME motif search tool (http://meme-suite.org/tools/meme) using default settings specifying a maximum of 10 motifs. The resulting motif architectures were aligned and visualized with the phylogenetic tree using TBtools-II (https://github.com/CJ-Chen/TBtools-II).

### HvAOC expression

Transcriptional expression data of *HvAOC* in various organs was obtained from EoRNA (https://ics.hutton.ac.uk/eorna/index.html) and BAR-ePlant Barley V3 (https://bar.utoronto.ca/). Except for the expression data of anthers, which was derived from Golden Promise, all other data were obtained from MOREX cultivar. The shoot was sampled when it reached 10 cm in length, leaves were collected two months after sowing, and roots were sampled four weeks after sowing. Lemma, palea, and lodicule were collected six weeks after flowering, rachis was collected five weeks after flowering, and anthers were collected when they reached a length of 1.3–1.4 cm. Grains were collected at five time points after flowering and divided into three parts including maternal tissue, endosperm, and embryo.

### TUNEL assay

Paraffin-fixed caryopses at 6 and 10 DAP were selected for TUNEL assay using the Dead End Fluorometric TUNEL Kit (Promega, G3250) according to the manufacturer’s instructions. Briefly, the paraffin was removed by xylene treatment, and the sections were rehydrated with an ethanol series. After treatment with proteinase K in PBS, the sections were incubated at 37 °C for 60 min in the presence of rTdT. The TUNEL-positive control was set with DNase treatment. The green fluorescence of fluorescein (TUNEL signal) and red fluorescence of propidium iodide were analyzed at 480 nm/520 nm and 620nm/640 nm on the excitation/emission spectrum under the Zeiss ZT0 confocal microscope. For each genotype, three independent plants were selected and sections from three individual grains per plant were used for TUNEL detection.

### Genetic analyses

The T3 *hvaoc-n4* homozygous and Cas-9-free line was crossed with the BWNIL938 containing the *Zeo1.b* gain of function *HvAP2* allele (Houston *et al*., 2013). The F2 population was genotyped for *HvAOC* and *HvAP2* alleles. F3 seed was collected from each F2 individual. Selected F3 families were taken forward for genotyping, phenotyping and grain analyses.

### RNAseq

Caryopses of Golden Promise and *hvaoc-n4* at 6 and 12 DAP were used for RNAseq. RNA isolation was performed using RNA Isolation and Purification Kit (Qiagen), followed with RNA integrity confirmation by Bioanalyzer 2100 (Agilent Technologies). Library construction, sequencing, data pre-processing and differential expression analysis were performed as in Liu *et al*. (2022) with differentially expressed genes (DEGs) identified using a log2 fold-change cutoff of 0.5. GO enrichment analysis of the DEGs was performed on g:Profiler using a custom barley GO annotation as described in Liu *et al*. (2022). The enriched GO terms were summarised using ReviGo before the visualisation.

### Spatial transcriptome

Barley grains (cv. Golden Promise) at 7 DAP and 10 DAP were dissected from hull and snap frozen in a liquid nitrogen cooled isopentane bath for 1 min, subsequently embedded in Optimal Cutting Temperature (OCT, Tissue-Tek). The grains were sectioned on a cryostat (Leica Biosystems) to 12 μm thick at –20°C. The sections were carefully place on a prechilled Tissue Optimization (TO) or Gene Expression (GE) slide from Visium Tissue Optimization Slide and Reagent Kit (10× Genomics, 1000193) and Visium Gene Expression Slide and Reagent Kit (10× Genomics, 1000187), respectively. As described by Peirats-Llobet *et al*. (2023), the sections were stained, imaged and incubated in pre-permeabilization solution at 37°C for 30 min. Following the Gene Expression workflow, cDNA from capture areas were synthesized and libraries were generated as described in 10× Genomics User guide (VisiumSpatialGeneExpression_UserGuide_RevE). Libraries were sequenced on an Illumina NextSeq 550 platform at Novogene (Australia). Raw reads, bright field image alignments and the latest barley reference genome (https://plants.ensembl.org/Hordeum_vulgare/Info/Index) were prepared as inputs for Space Ranger pipeline (https://www.10xgenomics.com/software) to generate transcript-spot matrices. Loupe Browser (v. 8.0, 10x Genomics) was used for gene expression visualization.

### Statistics

All statistical analyses were performed using IBM SPSS Statistics 26 (IBM Corp., Armonk, NY, USA). Depending on the experimental design, two-tailed Student’s t-tests, one-way ANOVA, and Tukey’s HSD post hoc tests were applied as indicated in the figure legends. Data visualisation was carried out using Origin 2021 and R, and final figure assembly was completed in Adobe Illustrator 2021. Measurements were obtained from at least three independent biological replicates, as specified in the corresponding figures, and no sample was measured more than once.

## Results

### HvAOC suppresses vegetative growth and developmental rate in barley

To explore the roles of JA in barley, we focused on the barley *ALLENE OXIDASE CYCLASE* (*AOC*) gene. Phylogenetic analyses confirmed that AOC is encoded by a single gene copy in many Poaceae, including barley (Fig S1a). Evaluation of conserved motifs across land plants (Fig S1b) suggested that AOC motif structure is largely conserved across grasses while haplotype analyses of *HvAOC* over diverse barley germplasm, including cultivated, landrace and wild barleys revealed 18 haplotypes (Fig S2). Of these, five HPs included non-synonymous SNPs (HPs_1, 2, 8, 9 and 15) and consisted entirely of wild barleys while cultivated HPs were represented in seven HPs (HPs 3,7,11, 13, 14, 15) with three exclusive to cultivated germplasm (HP 11, 14, 16) (Fig S2), which together suggest that *HvAOC* likely shows no functional variation across cultivated barley. Publicly available expression data mined from EoRNA (Milne *et al*., 2021) confirmed that *HvAOC* is expressed widely across different tissues (Fig S3). As *HvAOC* is a single copy gene whose ortholog encodes a biosynthetic enzyme early in the JA metabolic pathway and given its probable functional conservation, we selected *HvAOC* for gene editing to learn more about role(s) of JA in barley growth and development.

We edited *HvAOC* in the Golden Promise cultivar using the cloning strategy and transgenic protocol described in (Lawrenson *et al*., 2015; Lawrenson & Harwood, 2018) and screened the T2 generation for gene edits and identify plants lacking the *Cas9* transgene. We selected two putative impaired function alleles for further characterisation, *hvaoc-n4*, where an insertion and deletion are predicted to cause a frame-shift and early stop codon, and *hvaoc-n7*, where a 1-bp insertion similarly should cause a frame-shift and early stop codon (Fig 1a). Both lines were *Cas9*-free and propagated to the T4 generation for phenotypic comparison with parent cultivar Golden Promise.

**Figure 1.**
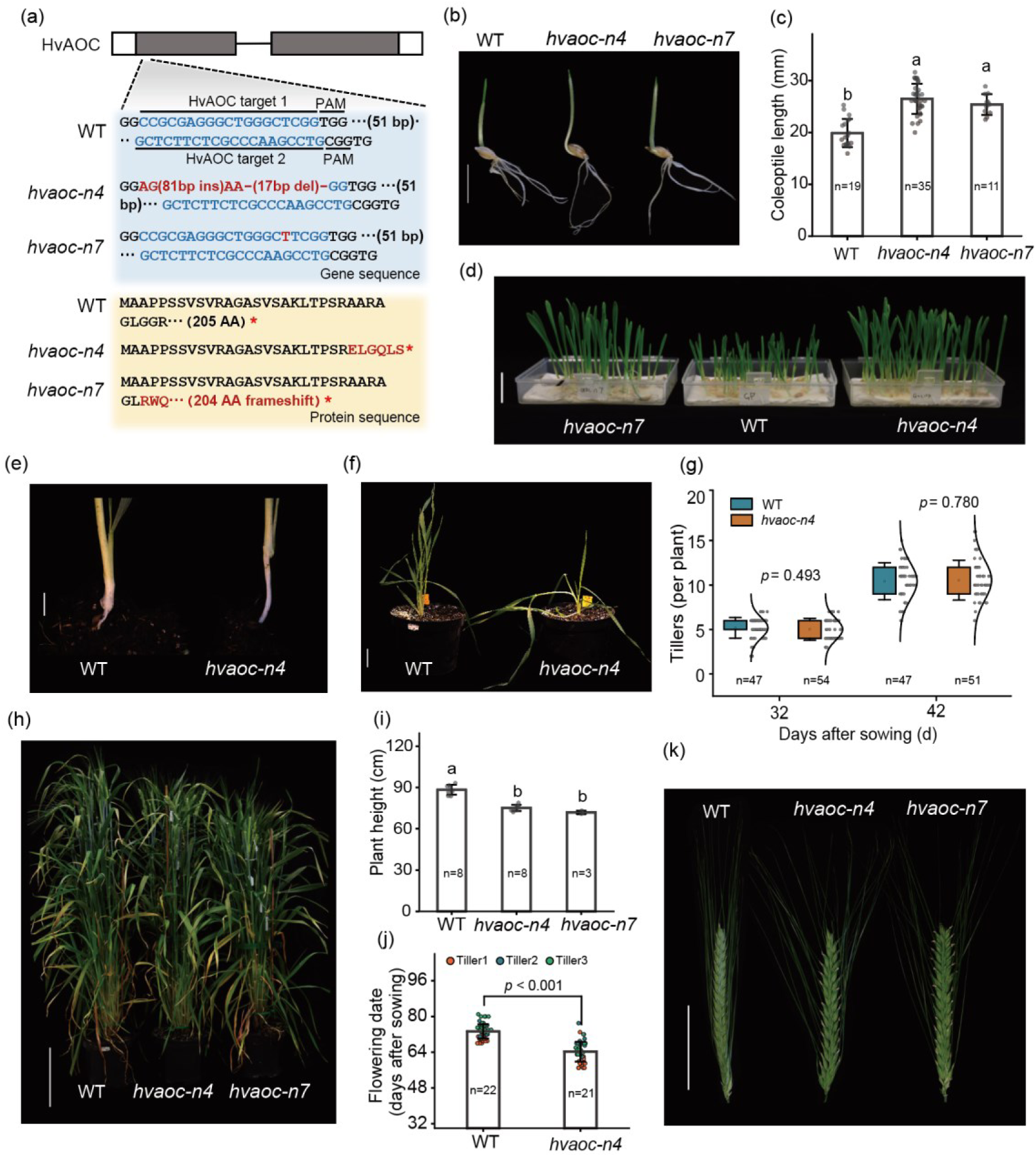
Generation and general phenotypes of *hvaoc* mutants. (a) *HvAOC* gene model with 5’ region of exon1 magnified in the inset to show the Golden Promise (WT) sequences targeted by the guides (blue shaded box) and corresponding protein sequence (yellow shaded box). Edited regions and predicted changes in protein sequence shown in red. (b-c) WT, *hvaoc-n4* and *hvaoc-n7* seedlings 5 days after sowing (DAS). (b), Scale bar, 1.5 cm. (c) Coleoptile length of WT, *hvaoc-n4* and *hvaoc-n7* seedlings 5 DAS. (d) WT, *hvaoc-n4* and *hvaoc-n7* seedlings at 8 DAS. Scale bar, 2 cm. (e-f) Subcrown internode of WT and *hvaoc-n4* at 17 DAS. Scale bar, 1 cm. (f) Growth habit of WT and *hvaoc-n4* plants at 32 DAS. (g) Tiller number of WT and *hvaoc-n4* plants 32 and 48 DAS. (h) Mature plants of WT, *hvaoc-n4* and *hvaoc-n7*. Scale bar, 20 cm. (i) Plant height of WT, *hvaoc-n4* and *hvaoc-n7* mature plants. (j) Flowering time of the three tallest tillers in WT, *hvaoc-n4* and *hvaoc-n7*. (k) Mature spikes of WT, *hvaoc-n4* and *hvaoc-n7* at 12 days after anthesis. Scale bar, 5 cm. Gray dots in (c), (g) (i) and (j) represent individual biological replicates (n = 3-54). Bars on the top of the column in (c), (i) and (j)represent standard deviation and different letters in (c) and (i) indicate significant differences (*P* < 0.05, one-way ANOVA, Tukey’s HSD test). Data in (g) and (j) were tested using Student’s t-test.

Germination rate was equivalent across Golden Promise, *hvaoc-n4* and *hvaoc-n7* (Fig S4); however, *hvaoc-n4* and *hvaoc-n7* seedlings had an 33% and 27% increased coleoptile length (Fig 1b, c), respectively, with seedlings showing clear increases in growth by 8 days after sowing (DAS, Fig 1d) compared to Golden Promise. We also noted longer subcrown internodes in *hvaoc-n4* compared to Golden Promise (Fig 1e). By 32 DAS, *hvaoc-n4* plants displayed a more prostrate growth habit compared to Golden Promise plants but no change in tiller number at either 32 DAS or 48 DAS (Fig 1f, g). Compared to Golden Promise, *hvaoc-n4* and *hvaoc-n7* were 15% and 18.7% shorter at maturity, respectively, (Fig 1h, i) and *hvaoc-n4* flowered 9.1 days earlier based on pollen dehiscence (Fig 1j). We detected no change in spike length, grain number and grain setting percentage per spike across the genotypes (Fig 1k, Fig S5). Taken together, loss of function *hvaoc* alleles lengthened coleoptiles, accelerated early seedling growth and flowered earlier compared to Golden Promise, suggesting a role for HvAOC in repressing early vegetative organ growth and stem elongation in barley.

### HvAOC represses caryopsis growth independently from the lemma

Grain on maturing *hvaoc-n4* and *hvaoc-n7* spikes appeared more prominent with lemmas splayed open compared to Golden Promise spikes (Fig 1k). Measuring grain parameters revealed that compared to Golden Promise, *hvaoc-n4* and *hvaoc-n7* grain was 13.6% and 13.3% longer and 14.4% and 5.3% thicker, respectively (Fig 2a-e), while lemma length remained unchanged across genotypes (Fig 2f), such that *hvaoc* grain bulged out to prise open the lemma, generating a more open architecture. Accordingly, while Golden Promise grain length matched lemma length, *hvaoc-n4* and *hvaoc-n7* grain exceeded the end of the lemma by 1.6mm and 1.2 mm, respectively. Uncoupling of lemma and grain length suggests that *HvAOC* may regulate grain dimensions independently from the lemma, in contrast to many other genes reported to control grain parameters. Grain weight was also altered with 11.3% and 8.7% heavier grain in *hvaoc-n4* and *hvaoc-n7* compared to Golden Promise, respectively (Fig 2g). Taken together, our data show that HvAOC is a major regulator of grain parameters, limiting grain length, depth and weight but not influencing lemma and palea elongation.

**Figure 2.**
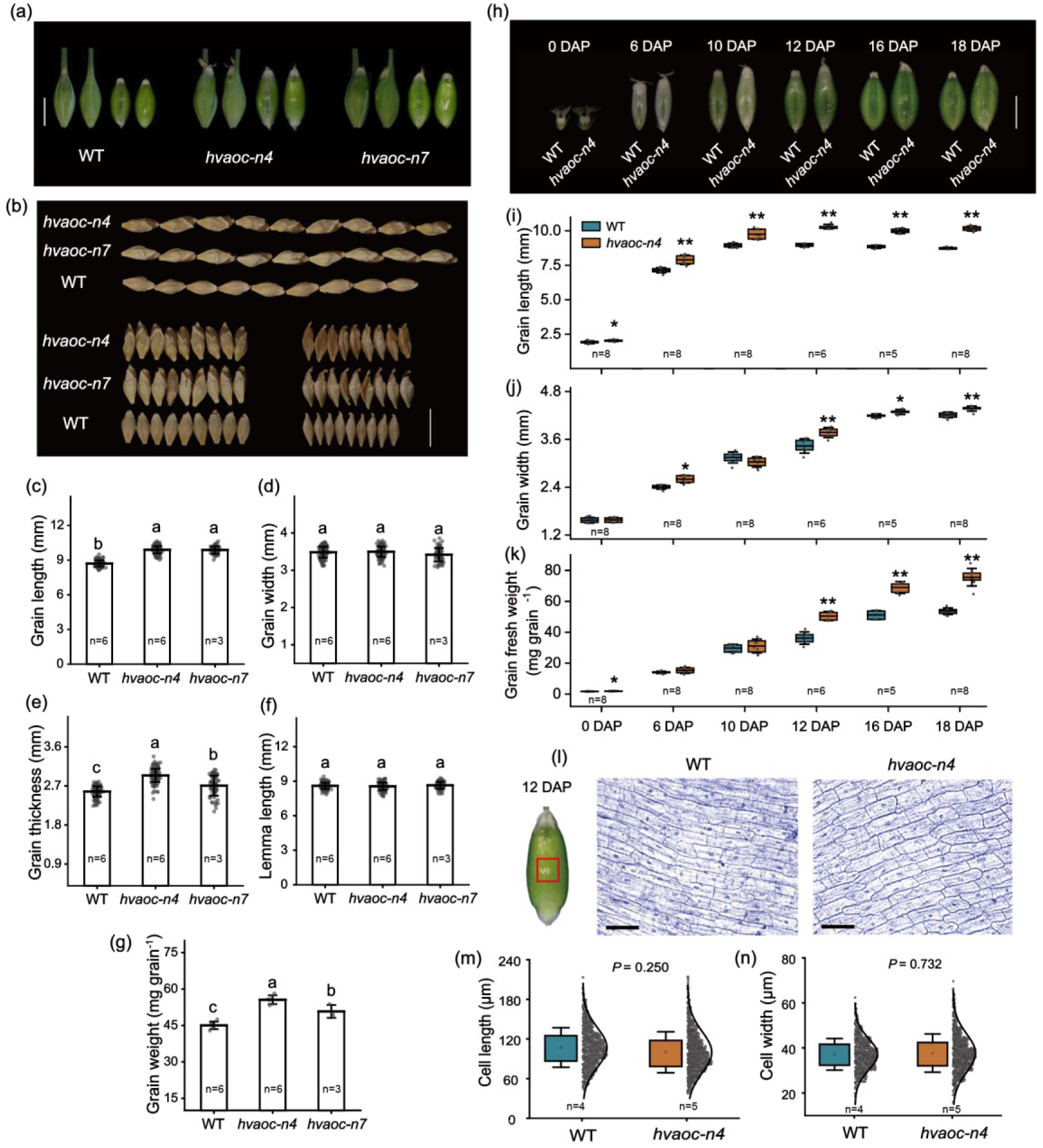
*HvAOC* controls grain parameters and development. (a) Caryopses removed from WT, *hvaoc-n4* and *hvaoc-n7* spikes 12 days after pollination (DAP), pictured with and without the lemma. Scale bar, 0.5 cm. (b) Mature grain of WT, *hvaoc-n4* and *hvaoc-n7.* The two sets are arranged differently to highlight changes in different dimensions: the top set is aligned to compare grain width, whereas the bottom set is aligned to compare grain thickness. Scale bar, 1 cm. (c-g) Comparisons of mature grain length (c), grain width (d), grain thickness (e), lemma length (f) and grain weight (g) between WT, *hvaoc-n4* and *hvaoc-n7*. The gray dots in (c)-(f) represent single grain measurement (24 measurements per individual) from at least three biological replicates, while the dots in (g) represent individual biological replicates. Bars on the top of the column represent standard deviation and different letters in (c)-(g) indicate significant differences (*p* < 0.05, one-way ANOVA, Tukey’s HSD test, n≥3). (h) Caryopsis development in WT and *hvaoc-n4* from 0 DAP to 18 DAP. Scale bar, 0.5 cm. (i-k) Comparisons of caryopsis length (i) width (j) and weight (k) between WT and *hvaoc-n4* from 0 DAP to 18 DAP. n ≥ 5 individuals per genotype. Data were tested using Student’s t-test, * and ** indicate significant difference at the 0.05 and 0.01 level, respectively. (l) Comparison of cell size in the ventral pericarp of WT and *hvaoc-n4* at 12 DAP. Representative photo on left showing the ventral region analyzed (red box). Centre and right panels show pericarp stained with Toluidine Blue. Scale bar, 100 μm. (m-n) Pericarp cell length and width. n ≥ 4 individuals per genotype, with 3 grain images per individual and ≥35 cell measurements per image. Data tested using Student’s t-test. Gray dots in (m) and (n) indicate one cell size measurement.

To explore the origin of grain differences, we tracked caryopsis development in *hvaoc-n4* and Golden Promise from 0 to 18 days after pollination (DAP) (Fig 2h-k). Compared to Golden Promise, *hvaoc-n4* caryopses were longer from 6 DAP onwards (Fig 2i) while grain width and fresh weight increased compared to Golden Promise at earlier and later stages (Fig 2j, k). Pericarp cell length and width in 12 DAP caryopses showed no difference between Golden Promise and *hvaoc-n4* suggesting that increased caryopsis length may represent increased cell number (Fig 2l-n).

### Phenotypes of *hvaoc* likely result from JA deficit

Given the putative role of HvAOC in JA biosynthesis, we hypothesized that *hvaoc* mutant phenotypes reflect a JA deficit. To test this, we treated both Golden Promise and the *hvaoc-n4* mutant with methyl jasmonate (MeJA) or mock solution and then looked for evidence of phenotypic rescue. While Golden Promise and *hvaoc-n4* developed shorter coleoptiles in response to 0.1 mM and 0.25 mM MeJA, these lengths were equivalent between Golden Promise and *hvaoc-n4*, suggesting that MeJA completely rescued the *hvaoc-n4* long coleoptile phenotype (Fig 3a, b). We also examined reproductive tissues treated with MeJA. Treatment of *hvaoc-n4* plants with 1 mM MeJA resulted in wild-type looking spikes without ‘prominent’ grain exceeding lemma boundaries (Fig 3c, d) and reduced grain length and thickness in both Golden Promise and *hvaoc-n4* (Fig 3e-g). The length and thickness of MeJA-treated *hvaoc-n4* grain was equivalent to Golden Promise under mock treatment and removed genotype differences in grain width (Fig 3f-h). Lemmas did not respond to MeJA treatment in either genotype (Fig 3i). Treatment with 1mM MeJA also reversed increased fresh caryopsis weight at 12 DAP and final grain weight in *hvaoc-n4* (Fig 3j, k). Altogether, *hvaoc-n4* mutant vegetative and reproductive phenotypes resembled Golden Promise phenotypes following treatment with MeJA, indicating that coleoptile and grain phenotype were associated with JA levels and loss of HvAOC function may lead to phenotypes in *hvaoc-n4* due to a deficit in JA.

**Figure 3.**
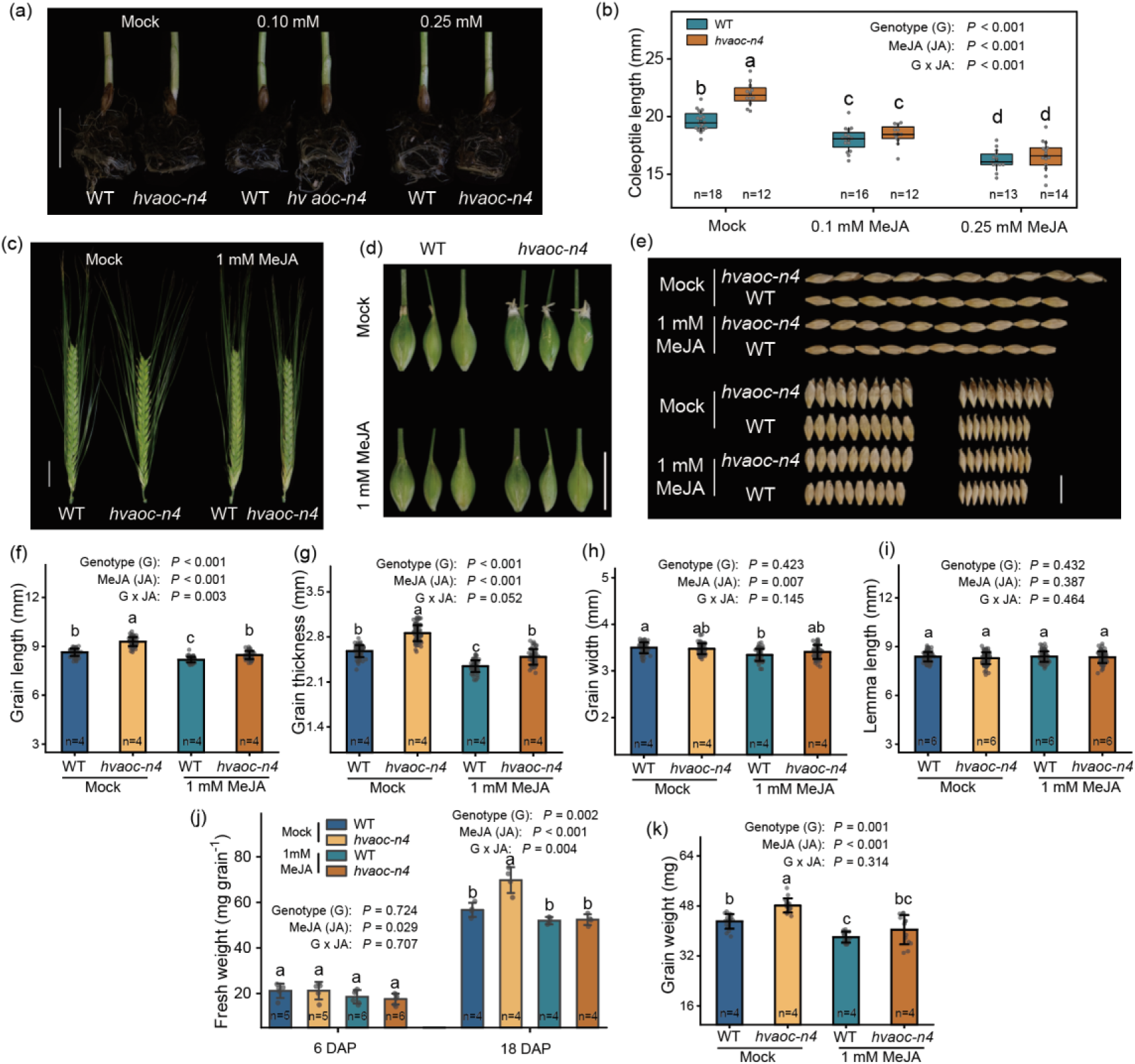
MeJA treatment on coleoptile and grain parameters in WT and *hvaoc-n4*. (a-b) Representative images of coleoptile length in WT and *hvaoc-n4* after mock and MeJA treatment. Scale bar, 2 cm. (b) quantification coleoptile length in WT and *hvaoc-n4* after mock and MeJA treatment. (c-d) Spike and grain morphology of WT and *hvaoc-n4* at 12 days after pollination (DAP) following mock or MeJA treatment. Scale bar, 2 cm. (e) Representative images of mature grains from WT and *hvaoc-n4* treated with mock or MeJA. Scale bar, 1 cm. Grain fresh weight at 6 DAP and 18 DAP under mock and MeJA treatments. (f) (g–k) Comparison of grain length (g), grain width (h), grain thickness (i), grain weight (j) and lemma length (k), in WT and *hvaoc-n4* under mock or MeJA treatments. Gray dots in (b) and (e) represent biological replicates, while gray dots in (g)-(k) represent one grain measurement taken from n ≥ 4 biological replicates. Different letters in (b), (e) and (g)-(k) indicate significant differences (*P* < 0.05, Two-way ANOVA, Tukey’s HSD test, n ≥ 4).

### HvAOC regulates grain size parentally and independently of HvAP2

Our previous work showed that HvAP2 restricts grain length which was associated with HvAP2 promotion of JA-associated gene expression (Shoesmith *et al*., 2021). To examine genetic interactions between *HvAP2* and *HvAOC*, we crossed the BwNIL938 (Bowman near isogenic line 938) containing *Zeo1.b*, a *miR172*-resistant gain of function *HvAP2* allele (Houston *et al*., 2013) with Cas9-free *hvaoc-n4* T3 plants. The Golden Promise cultivar background for the *hvaoc-n4* edits contains the mild *Zeo2* allele (Houston *et al*., 2013). We genotyped F2 individuals for *Zeo* and *HvAOC* alleles (χ2 =4.63, n = 64, P = 0.80) and selected those fixed for either *Zeo1.b* or *Zeo2* allele and heterozygous at the *HvAOC* locus (i.e. segregating for Bowman *HvAOC* and the gene-edited *hvaoc-n4* alleles). We planted and genotyped F3 families from each selected F2. Each F3 family contains a mosaic Bowman and Golden Promise backgrounds. Within each family (*Zeo2*/ *Zeo2 hvaoc-n4/ hvaoc-n4* F3, χ2 = 0.40, n = 63, P = 0.82; *Zeo1.b*/ *Zeo1.b hvaoc-n4*/ *hvaoc-n4* F3, χ2 = 0.37, n = 60, P = 0.83; Table S4), we identified and phenotyped *HvAOC*/ *HvAOC*, *HvAOC*/ *hvaoc-n4* and *hvaoc-n4*/ *hvaoc-n4* siblings. We first compared *HvAOC*/ *HvAOC* and *hvaoc-n4*/ *hvaoc-n4* siblings fixed for either the *Zeo2* or *Zeo1.b* allele. *Zeo1.b* causes shorter internodes in the stem and spike (Houston *et al*., 2013; Patil *et al*., 2019). We detected no difference in spike density in the *Zeo1.b*/ *Zeo1.b* siblings regardless of *HvAOC* or *hvaoc-n4* allele (Fig 4a, c), suggesting that HvAOC does not influence HvAP2 suppression of rachis (spike axis) internode length. However, *hvaoc-n4*/ *hvaoc-n4* siblings combined with either *Zeo2* or *Zeo1.b* alleles showed slight increases in plant height (Fig S6), suggesting that *hvaoc-n4* can partially compensate for *Zeo*-associated decreases in stem elongation. Grain formed on *Zeo2*/ *Zeo2 hvaoc-n4/ hvaoc-n4* and *Zeo1.b*/ *Zeo1.b hvaoc-n4/ hvaoc-n4* spikes was visible compared to other siblings (Fig 4b) consistent with caryopses exceeding the lemma boundaries. Measuring grain length confirmed that *Zeo2*/ *Zeo2 hvaoc-n4/ hvaoc-n4* and *Zeo1.b*/ *Zeo1.b hvaoc-n4/ hvaoc-n4* developed longer grain than siblings with two copies of *HvAOC* (Fig 4d) The type of *Zeo* allele made no difference to grain length in *HvAOC*/ *HvAOC* individuals; however, grain from *Zeo1.b*/ *Zeo1.b hvaoc-n4/ hvaoc-n4* plants were shorter compared to *Zeo2*/ *Zeo2 hvaoc-n4*/ *hvaoc-n4* indicating that *Zeo1.b* can partially compensate for the grain-lengthening effect of *hvaoc-n4* (Fig 4d). Genotypes showed no differences in grain width or thickness (Fig 4e, f) or grain weight, except for *Zeo2*/ *Zeo2 hvaoc-n4*/ *hvaoc-n4* plants which yielded heavier grain (Fig 4h). Interestingly, *Zeo2*/ *Zeo2 hvaoc-n4*/ *hvaoc-n4* florets had longer lemmas compared to other genotypes (Fig 4g). Since *hvaoc-n4* did not change lemma length in the Golden Promise background, increased lemma length in *Zeo2*/ *Zeo2 hvaoc-n4*/ *hvaoc-n4* plants may reflect interactions between parental backgrounds or that HvAOC may influence lemma length in a *Zeo* allele-dependant manner. Taken together, gain of HvAP2 function partially interfered with the *hvaoc-n4* ability to increase grain length and weight, suggesting that HvAP2 restriction of grain length may depend on adequate activity of HvAOC, and therefore, the JA pathway.

**Figure 4.**
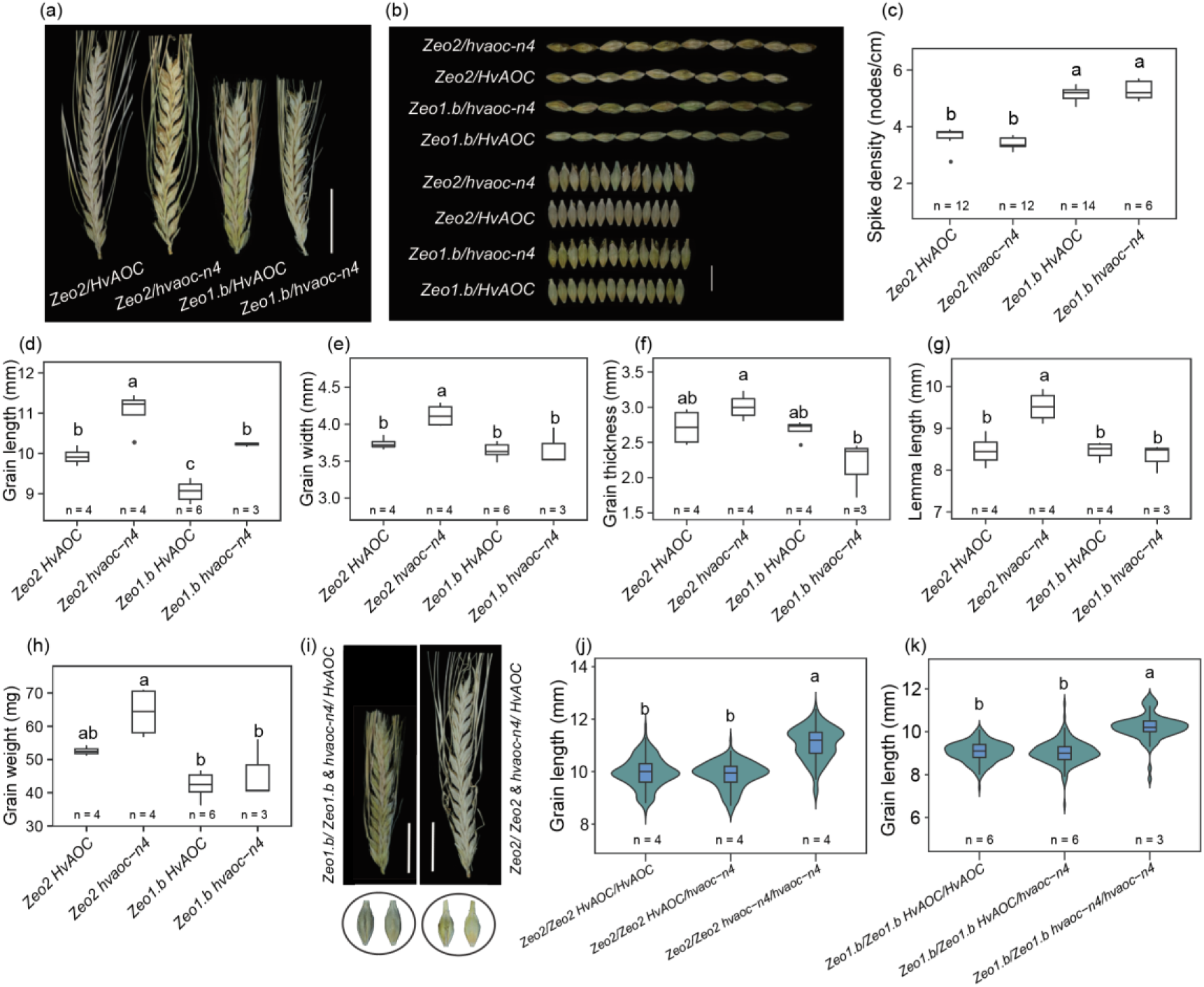
Grain traits and lemma length in *Zeo1.b* x *hvaoc-n4* F3 siblings. (a-b) Mature spike and grain morphology of *Zeo2 HvAOC* (*Zeo2*/ *Zeo2 HvAOC/ HvAOC*), *Zeo2 hvaoc-n4* (*Zeo2*/ *Zeo2 hvaoc-n4/ hvaoc-n4*, *Zeo1.b hvaoc-n4* (*Zeo1.b*/ *Zeo1.b hvaoc-n4/ hvaoc-n4*) and *Zeo1.b HvAOC* (*Zeo1.b*/*Zeo1.b HvAOC/ HvAOC*). Scale bar in (a), 3 cm. Scale bar in (b), 10 mm (c-h) Quantification of spike and grain morphological traits, including spike density(c), grain length, (d), grain width (e), grain thickness (f), grain weight (g) and lemma length (h). The lower and upper box edges represent the first and third quartiles, the horizontal lines within box indicate the median, and the lower and upper whiskers denote the minimal and maximal values minimal and maximal values within 1.5* interquartile range, respectively, while points indicate outliers beyond this range. *N* ≥ 3 per genotype. (i) Spike and grain morphology of *Zeo2*/ *Zeo2 hvaoc-n4/ HvAOC* and *Zeo1.b*/ *Zeo1.b hvaoc-n4/ HvAOC* at mature stage. Scale bar, 3 cm. (j-k) grain length of *HvAOC / HvAOC, hvaoc-n4/ HvAOC* and *hvaoc-n4/ hvaoc-n4* both with *Zeo2*/ *Zeo2* and *Zeo1.b*/ *Zeo1.b* at mature stage. The lower and upper box edges within violin plot represent the first and third quartiles, the horizontal lines within box indicate the median, and the lower and upper whiskers denote the minimal and maximal values minimal and maximal values within 1.5* interquartile range, respectively. n ≥ 3 per genotype, with 178, 176 and 119 grains measured for *Zeo2/Zeo2 HvAOC/HvAOC*, *Zeo2*/ *Zeo2 hvaoc-n4/ HvAOC1* and *Zeo2*/ *Zeo2 hvaoc-n4/ hvaoc-n4*, 190, 186 and 86 grains measured for *Zeo1.b*/ *Zeo1.b HvAOC/HvAOC*, *Zeo1.b*/ *Zeo1.b hvaoc-n4/ HvAOC1* and *Zeo1.b*/ *Zeo1.b hvaoc-n4/ hvaoc-n4*, respectively. Different lowercase letters in (c-h) and (j-k) indicate significant differences (*P* < 0.05, One-way ANOVA, Tukey’s HSD test).

We also examined grain parameters in F3 plants genotyped as *HvAOC*/ *hvaoc-n4* and fixed for either *Zeo* allele; all grain from these plants will have maternal tissues with the *HvAOC*/ *hvaoc-n4* genotype but contain embryos with these genotype distributions: 25% *hvaoc-n4*/ *hvaoc-n4*, 50% *HvAOC*/ *hvaoc-n4*, and 25% *HvAOC*/ *HvAOC*. Grain from *HvAOC*/ *hvaoc-n4* siblings with either *Zeo* allele, did not show visible differences or different populations of grain length compared to grain from *HvAOC*/ *HvAOC* siblings (Fig 4i-j), while grain from *hvaoc-n4*/ *hvaoc-n4* siblings showed longer grain as expected (Fig 4d, j-k). Therefore, changes in grain length due to *hvaoc-n4*/ *hvaoc-n4* are parentally-derived, suggesting that HvAOC function in parental tissues regulates grain dimensions.

### HvAOC controls maternal tissue differentiation and degradation

Differences in grain parameters between Golden Promise and *hvaoc-n4* alleles emerged early during grain development (Fig 2). To understand more about the relationship between HvAOC function and grain development, we examined *HvAOC* expression in the BAR-ePlant Barley V3 RNAseq dataset from developing caryopses (https://bar.utoronto.ca/; Kovacik *et al*., 2024), which revealed higher expression of *HvAOC* in maternal versus filial grain tissues (Fig 5a). We next examined *HvAOC* expression in caryopses using Golden Promise spatial transcriptome at 10 DPA (Fig 5b). *HvAOC* transcripts were detected across the grain, and noticeably enriched in vascular bundles, flanking vascular bundles and the nucellar projection (Fig 5b), tissues with important roles in parental to filial nutrient transfer (Radchuk *et al*., 2023).

**Figure 5.**
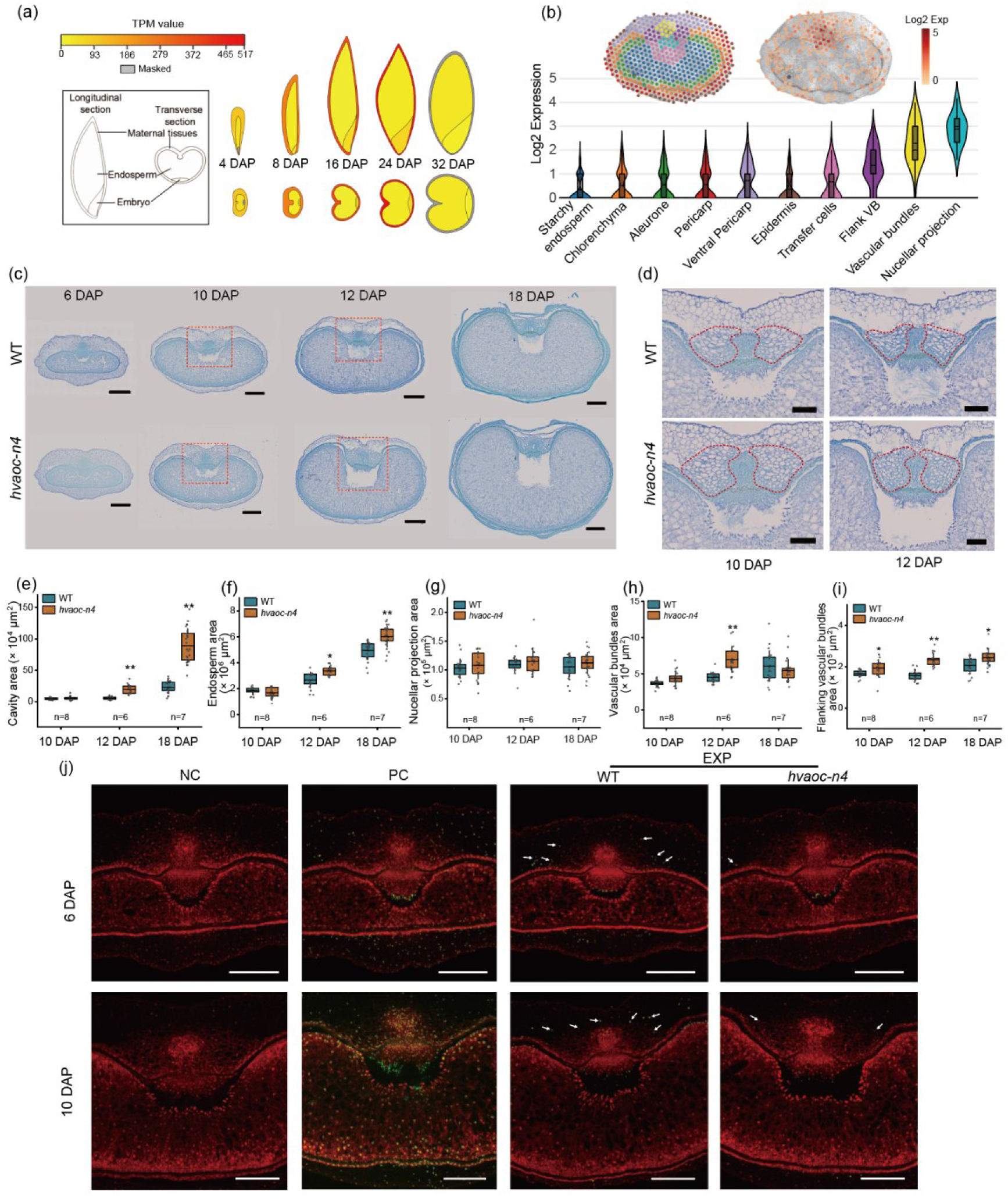
*HvAOC* expression and influence on grain tissue development. (a) Transcript levels of *HvAOC* detected in parental tissues, endosperm and embryo at different days after pollination (DAP), visualized from BAR-ePlant Barley V3 (https://bar.utoronto.ca/). Colours represent mean TPM values (n = 3) according to the heat map scale. Gray color of parental tissues at 32 DAP indicates an expression level over 100% of their standard deviation. TPM, transcripts per million. (b) Spatial expression pattern of *HvAOC* at 10 DAP in different grain tissues revealed by spatial transcriptomics. Violin plots show expression in spots from different grain tissues. Grain image on left shows tissues and image on right shows *HvAOC* distribution on spots. (c-d) Grain transverse sections of Golden Promise wild-type (WT) and *hvaoc-n4* at 6, 10, 12 and 18 DAP stained with toluidine blue. Red box indicates region at higher magnification in (d), scale bars, 500 μm. (d) Magnified images of the nucellar projection (NP), vascular bundles (VB), flanking vascular bundles (FVB) and cavity from sections shown in (c). Red dashed lines indicate the FVB region, scale bars, 200 μm. (e–i) Image J quantification cavity (e), starchy endosperm (f), nucellar projection (g), vascular bundles (h), and flanking vascular bundles (i) area. Data were tested using Student’s t-test, * and ** indicate *P* < 0.05 and *P* < 0.01. *N* ≥ 6 per genotype, with 3 grain sections measured per individual as technical replicates. Gray dots in (e-i) represent one tissue area measurement. (j) Detection of programmed cell death by TUNEL assay in 6 and 10 DAP caryopsis sections. Green fluorescence indicates TUNEL-positive nuclei; white arrows mark TUNEL signals at the edge of the FVB in WT. Red fluorescence shows propidium iodide (PI) counterstaining. NC, negative control; PC, positive control; EXP, experimental treatment. Scale bars, 400 μm.

Examining grain anatomy in Golden Promise compared *hvaoc-n4* mutant by semi-thin sections revealed profound differences. Grain maturation in cereals is marked by the appearance of a so-called cavity, associated with programmed cell death (PCD) of maternal tissues, especially the nucellar projection (Dominguez & Cejudo, 2014; Radchuk *et al*., 2006; Shoesmith *et al*., 2021). The cavity expanded progressively from 6 to 18 DAP in both genotypes (Fig 5c); however, cavity area of 12 and 18 DAP caryopses was 3.5- and 3.9-fold larger in *hvaoc-n4* caryopses than Golden Promise, respectively (Fig 5c, e). Similarly, at 18 DAP, endosperm area was 21.6% greater in *hvaoc-n4* caryopses than Golden Promise (Fig 5c, f). We did not observe any significant differences in NP area between genotypes in 10, 12 or 18 DAP caryopses (Fig 5g). By contrast, the vascular bundle (VB) area increased 55.3% in *hvaoc-n4* 12 DAP caryopses compared to equivalently staged Golden Promise (Fig 5h). Notably, flanking vascular bundle (FVB) area at 10, 12 and 18 DAP increased by 16.9%, 48.6% and 17.7% in *hvaoc-n4* caryopses compared to Golden Promise, respectively (Fig 5d, i). Altogether, loss of HvAOC enhanced the area and integrity of the VB and FVBs as well as cavity expansion, putatively increasing assimilate allocation into the filial tissues. Since the endosperm cavity appears concomitant with maternal tissue PCD, we asked whether *HvAOC* regulates PCD in developing grain. We used TUNEL assays which label DNA in apoptotic cells with fluorescein-12-dUTP. We first observed TUNEL signal positive nuclei in Golden Promise caryopses at 6 DAP, mostly localized to regions laterally adjacent to the FVBs; by 10 DAP, the TUNEL signal localised at the upper FVB margins (Fig 5j). In contrast, *hvaoc-n4* 6 DAP caryopses exhibited 64.9% fewer TUNEL signal spots adjacent to the FVB compared to Golden Promise (Table S5) while TUNEL-positive nuclei persisted in the surrounding pericarp at 10 DAP. Although the distribution of TUNEL-positive nuclei in the NP region appeared similar between *hvaoc-n4* and WT by 10 DAP, Golden Promise showed more TUNEL signal spots (Fig 5j, Table S5). Fewer TUNEL-positive nuclei in the VB and FVB of *hvaoc-n4* compared to Golden Promise suggests less degeneration in these tissues which marries well with their larger area in grain sections. Taken together, *HvAOC*, and accordingly JA, may be required for timely PCD in maternal vascular tissues.

### Loss of HvAOC function influences gene expression associated with sugar transport, endosperm development, cell death and defense signalling

To determine how loss of HvAOC function influences gene expression during grain development, we performed comparative RNAseq on 6 and 12 DAP caryopses harvested from Golden Promise and *hvaoc-n4* mutants. We detected 2055 and 406 differentially expressed genes (DEGs) between genotypes at 6 and 12 DAP, respectively, with 52 DEGs shared between stages (Fig 6a; Table S6). Plotting spatial patterning of DEGs revealed highest concentration in parental transfer tissues in 6 DAP grain but also DEGs in filial tissues such as the endosperm and transfer cells (Fig 6b). DEGS at 6 DAP were enriched for several Gene Ontogeny (GO) terms (Fig 6c, Table S7), such as seed developmental processes (GO:0009793), which included genes encoding Sugars Will Eventually be Exported Transporters (SWEETs)(Fig 6e): *hvaoc-n4* 6 DAP caryopses showed 4.6-fold increased *HvSWEET14a* (HORVU.MOREX.r3.1HG0010640) and 2.78-fold increased *HvSWEET14b* (HORVU.MOREX.r3.6HG0540540) expression, the latter gene known to promote sucrose transport from maternal tissues into the grain (Radchuk et al., 2023). Other GO enriched terms included primary meristem (GO:0010072) (TPLs and PHB) and inflorescence development (GO:0048281), including auxin transporter BIG (HORVU.MOREX.r3.5HG0469420) and ERECTA kinases (Fig 6e, Table S7). We also detected downregulated DEGs under enriched terms for starch catabolism (GO:0005983) and beta-glucan synthesis (GO:0006075). While not associated with an enriched GO-term, the barley gene (HORVU.MOREX.r3.6HG0624320) was strongly downregulated in *hvaoc-n4* and is predicted to encode a class I MADS-box-encoding gene orthologous to *OsMADS87* which is epigenetically repressed to promote timely endosperm cellularisation (Folsom *et al*., 2014; Chen *et al*., 2016) (Fig 6e, Table S7). By 12 DAP, enriched terms included sucrose synthesis (GO:0005986), representing several upregulated fructose-1,6-bisphosphatases, and gluconeogenesis (Fig 6d, Table S7). We also detected enrichment GO terms such as response to water (GO:0009414) and vacuolar processing (GO:0006624), which included genes encoding vacuolar processing enzymes which were downregulated in *hvaoc-n4*, such as *VPE2c* (HORVU.MOREX.r3.2HG0184580) downregulated 1.88-fold and *VPE2b* (HORVU.MOREX.r3.2HG0184570) downregulated 1.74-fold (Fig 6e), the latter gene previously shown as expressed in the NP and associated with PCD (Radchuk *et al*., 2011). Cereal endosperm, including that in barley, is O_2_-deficient (Rolletschek *et al*., 2004). Interestingly, the enriched term Response to hypoxia (GO:0071456) represented DEGs all upregulated in *hvaoc-n4* mutant (Table S7), while one of the most highly upregulated genes in *hvaoc-n4* at 12 DAP encodes a AP2-ERF transcription factor (HORVU.MOREX.r3.2HG0112960), member of a family known to mediate hypoxia-related gene expression in Arabidopsis (Yang *et al*., 2011). While no SWEET-encoding genes contributed to GO-enriched terms at 12 DAP, the *hvaoc-n4* mutant showed 1.5-fold upregulation of *HvSWEET14a* (HORVU.MOREX.r3.1HG0010640), similar to 6 DAP, and a 1.76-fold upregulation of *HvSWEET11b* (HORVU.MOREX.r3.7HG0684580; Table S7b), reported to be highly expressed in nucellar parental tissues and to transport sucrose and cytokinin and promote endosperm cell number (Radchuk *et al*., 2023). In addition to *HvSWEET14a*, the most highly upregulated DEGs shared between 6 and 12 DAP in *hvaoc-n4* mutants included Pathogenesis-related (PR) and TGA transcription factors, defense genes which could be related to changes in PCD and/or differences in cavity formation (Table S7c). Another shared DEG encodes gibberellin oxidase 20 (HORVU.MOREX.r3.4HG0331680), a key biosynthetic step in producing the growth-promoting gibberellin phytohormone (ref) that was upregulated 2.8-fold at 6 DAP and 3.2-fold a 12 DAP in *hvaoc-n4* mutants. Additionally, the barley ortholog (HORVU.MOREX.r3.2HG0174520) of the rice *DWARF4* gene, which encodes a cytochrome P450 essential for brassinosteroid biosynthesis and which promotes grain length (Tanabe *et al*., 2005), was 17-fold elevated in the mutant. Taken together, GO-enrichment and individual genes differentially expressed in *hvaoc-n4* caryopses, especially the *HvSWEET*, *HvVPE*, phytohormone and defense genes, suggest changes in parental tissue development and increased nutrient flow which may underlie formation of enlarged endosperm in *hvaoc-n4*.

**Figure 6.**
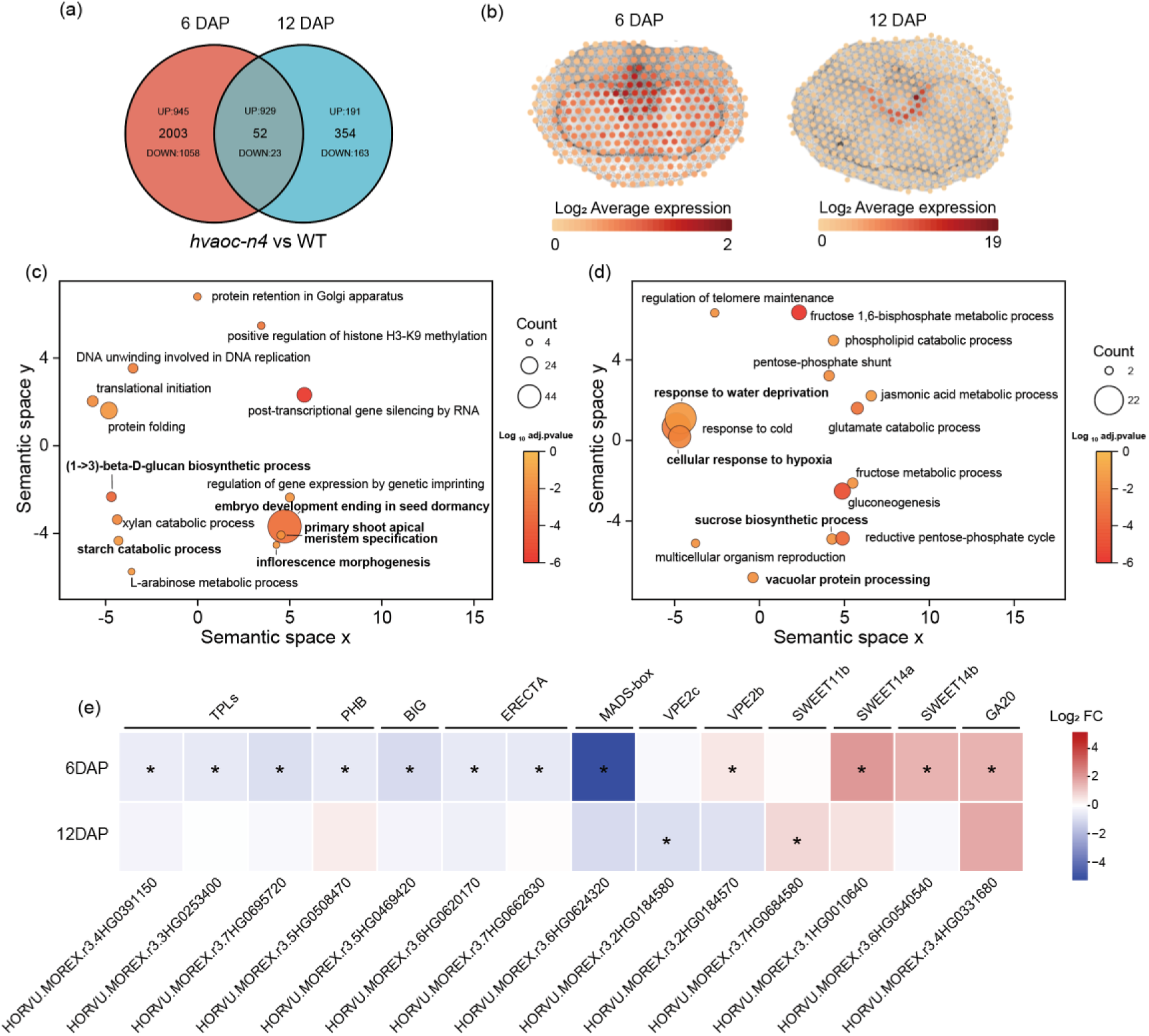
Transcriptomic alterations in the hvaoc-n4 mutant during early grain development. (a) Venn diagram showing differentially expressed genes (DEGs) in *hvaoc-n4* compared with wild type (WT) at 6 and 12 days after pollination (DAP). (b) Expression landscape of DEGs showing the average expression intensity of DEGs at 6 and 12 DAP mapped to 7 DPA and 10 DPA grain. (c-d) Semantic similarities between significantly enriched Gene Ontology (GO) categories in DEGs in *hvaoc-n4* compared with WT at 6 and 12 DAP. Bubbles represent GO terms whose closeness corresponds with semantic similarity. Bubble colours represent p values and bubble sizes represent the number of DEGs carrying this term. (e) Heatmaps based on expression fold changes of representative DEGs related to grain development in *hvaoc-n4* compared with WT at 6 and 12 DAP. DEGs with expression |log_2_ fold change|≥0.5 and a Benjamini–Hochberg adjusted p-value ≤ 0.05 are highlighted by asterisks.

### JA biosynthetic and signaling pathway genes show distinctive expression in the grain spatial transcriptome

We also examined genes encoding all steps in the JA biosynthetic pathway across the spatial transcriptome at 7 DAP (Fig S7). Within multigene families, i.e. not *HvAOC*, selected individual genes showed distinctively higher expression compared to other members suggesting more important roles in JA production during grain development. Interesting, the barley orthologs of *LOX* and *AOS* genes involved in early steps in JA biosynthesis were enriched in maternal tissues: for instance, transcripts of *HvLOX* genes (HORVU.MOREX.r3.5HG0509200, HORVU.MOREX.r3.4HG0335790 and HORVU.MOREX.r3.6HG0614350) and the *HvAOS* genes (HORVU.MOREX.r3.4HG0394980 and HORVU.MOREX.r3.5HG0513120) appeared more frequent in the pericarp and vascular regions while *HvAOC* localized to pericarp vascular bundles and the nucellar projection (Fig 7, Fig S7). Interestingly, the most highly expressed *HvAOS* gene HORVU.MOREX.r3.4HG0394980 was downregulated in the *hvaoc-n4* mutant, as were several other biosynthetic genes (Fig S7) possibly suggesting negative feedback. In contrast, gene families associated with later steps in JA biosynthesis were more broadly expressed (Fig 7, Fig S7). Of particular interest were *HvOPR* genes, HORVU.MOREX.r3.7HG0726360, HORVU.MOREX.r3.4HG0331730 and HORVU.MOREX.r3.2HG0170590, which were most strongly expressed in vascular bundles, nucellar projection and filial transfer cells, respectively (Fig 7, Fig S7). We also examined genes encoding key signalling components including the COI1 receptor, JAZ inhibitors and TPL and NINJA co-factors. At 7 DAP, *HvCOI1* transcripts were clearly enriched in maternal tissues, while *JAZ* transcripts were similarly expressed in maternal tissues, especially the nucellar projection, but with some minor expression in filial tissues (Fig 7, Fig S8). On the other hand, *HvNINJA* transcripts were most highly expressed in transfer cells and moderately expressed in all tissues with the while *HvTPL*s were highly expressed in all tissues. At 10 DAP, JA biosynthesis genes were similarly expressed as in 7 DAP with early steps were still mostly limited to maternal tissues and later steps more broadly expression. However, while *HvCOI1* expression pattern seemed little changed at 10 DAP compared to 7 DAP, we detected a shift in the *HvJAZ* gene (HORVU.MOREX.r3.2HG0203510) which was little expressed at 7 DAP but strongly upregulated in all tissues at 10 DAP with highest expression in vascular bundles and the nucellar projection. We detected other less dramatic changes in the TPL and NINJA genes (Fig S8). Interestingly, we detected little expression and no tissue specificity for MYC2, the classic TF downstream of JA signalling, which was also absent from the DEGs.

**Figure 7.**
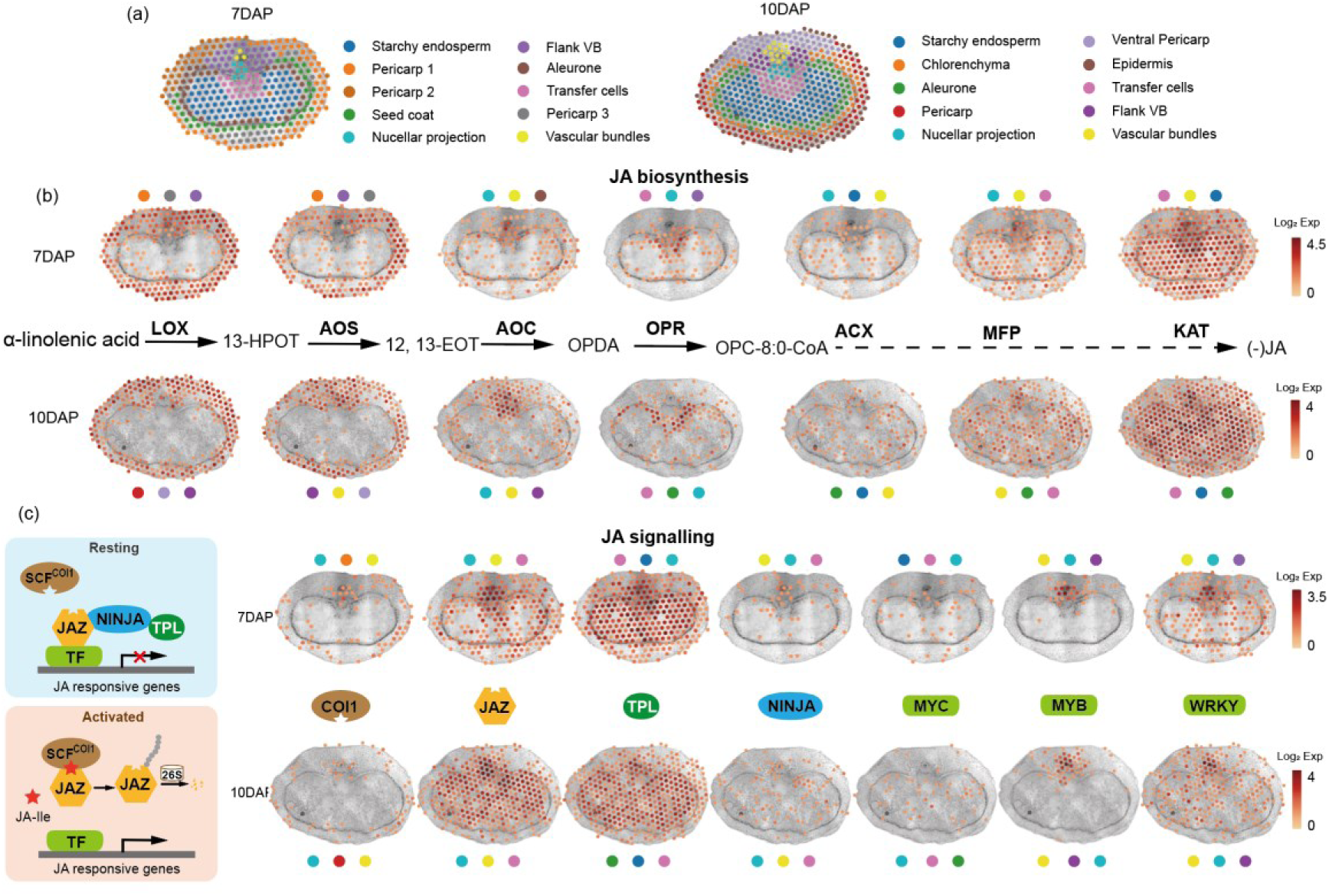
Spatial expression patterns of JA biosynthesis and signalling genes in barley grains at 7 and 10 DAP. (a) Tissue annotation of spatial transcriptomic sections from barley grains at 7 and 10 DAP. Tissues were generated using K-means pipeline and the cluster number was set at 10. (b) Spatial expression maps of key enzymes in JA biosynthetic pathway at 7 and 10 DAP. Genes involved in successive steps from α-linolenic acid to (–)-JA, including LOX, AOS, AOC, OPR, ACX, MFP, and KAT, are shown. The spatial heatmap show the results of highest expressed gene of each gene family based on the RNA-seq expression data. 13-HPOT, 13(S)-hydroperoxyoctadecatrienoic acid; 12,13-EOT, 12,13(S)-epoxyoctadecatrienoic acid; OPDA, (9S,13S)-12-oxo-phytodienoic acid; OPC 8:0-CoA, 3-oxo-2(cis-2ʹ-pentenyl)-cyclopentane-1-octanoic acid CoA. LOX, lipoxygenase; AOS, allene oxide synthase; AOC, allene oxide cyclase; OPR, OPDA reductase; ACX, acyl-CoA oxidase; MFP, multifunctional protein; KAT, 3-ketoacyl-CoA thiolase. (c) Schematic representation of the JA signalling pathway (left), illustrating the JAZ–NINJA–TPL repressor complex in the resting state and COI1-mediated JAZ degradation upon JA-Ile activation. Spatial expression maps (right) display the distribution of major JA signalling components at 7 and 10 DAP, including COI1, JAZ, TPL, NINJA and transcription factors (MYC, MYB and WRKY). Spatial heatmaps in (b) and (c) represent log₂-transformed UMl counts across tissue domains. Coloured circles above and below each heatmap in (b) and (c) denote the top three tissue domains with the highest expression for the corresponding gene. For key JA biosynthesis enzyme genes and signaling components, only the members of the gene family with the highest expression is used to generate the heatmap, the average expression heatmaps of whole gene family are shown in Figure S7 and Figure S8.

## Discussion

We tested our hypothesis that JA regulates vegetative and reproductive growth in barley by editing the *HvAOC* gene, a single copy gene encoding an enzyme acting early in JA biosynthesis. We characterised two Cas9-free lines with different predicted loss of function mutations in *HvAOC*. Our results suggest that HvAOC function is important to restrict coleoptile and early seedling growth and delays flowering based on time to anthesis (Fig 1). Two rice mutants with defective *AOC* alleles were reported to show enhanced coleoptile growth and earlier heading, suggesting these roles are conserved between rice and barley. However, the rice mutants also formed elongated floret hulls (Reimann *et al*., 2013) which we did not observe in *hvaoc* mutants in the Golden Promise background. Nonetheless, *hvaoc* mutants containing mixed Bowman and Golden Promise backgrounds developed longer lemmas compared to siblings with the *HvAOC* wild type allele (Fig 4), suggesting that other genes may interact with HvAOC to regulate this phenotype.

The visually striking splayed lemmas observed in *hvaoc* mutants derived from longer and thicker grain which prised open the floret hulls (Fig 1k, Fig 2a). Caryopses dimensions differed between Golden Promise and *hvaoc-n4* early in development, and corresponded to increased maternal tissues, especially of the vascular and flanking vascular bundles, as well as increased endosperm cavity and endosperm area (Fig 5). In Arabidopsis, JA signalling represses integument cell proliferation and ovule size (Hu *et al*., 2021). Mutant integument and pericarp layers must be larger as they enclose the larger seed, and as our data shows pericarp epidermis cells did not change in size, we suggest that larger tissues reflect increased cell proliferation. Our previous work showed that HvAP2 restricted pericarp cell number and limits integument layer number, as well as controlling the rate of maternal tissue degradation (Shoesmith *et al*., 2021). Thus, we speculate that *Zeo1.b* mitigates *hvaoc* grain lengthening in double mutants by partially suppressing enhanced proliferation in maternal layers due to the loss of HvAOC.

In rice, *coi* mutants and overexpression of the JA signalling repressor *OsJAZ11* increases grain width and weight (Mehra *et al*., 2022). Interestingly, in vitro assays found a direct interaction between OsJAZ11 and OsMADS29, a key regulator of NP differentiation and death, suggestive of a direct role for JA signalling in parental tissues, especially the NP. Our anatomical, TUNEL assay and transcriptomic results also support a role for JA in promoting maternal tissue degradation, potentially upstream of a JAZ11-MADS29 module. While parental death is important for filial tissue growth (Dominguez & Cejudo, 2011), increased persistence and size of flanking vascular bundles in *hvaoc* mutants correlated with increased cavity and endosperm area (Fig 5), as well as transcriptomic evidence for enhanced nutrient transport (Fig 6). As vascular tissues transport nutrients and sugar metabolites from parental reserves into grain (Olsen, 2004), we speculate that larger transport tissues and/or cavity volumes increase nutrient flow, consistent with previous work suggesting that enlarged cavities facilitate assimilate transport and enhance endosperm growth (Radchuk *et al*., 2021). Furthermore, since the endosperm itself constitutes an O_2_ diffusion barrier (Langer *et al*., 2023), we speculate that the larger endosperm volume and increased starch biosynthesis could increase the level of hypoxia, as reflected in the hypoxia-related DEGs upregulated *hvaoc*. Hypoxia is a state important for nutrient transit and starch metabolism (Borisjuk & Rolletschek, 2009) so increased endosperm volume may feed forward to further reduce O_2_ diffusion. A recent study in maize showed that a monocot-specific homolog of Inducer of CBF Expression 1 (ZmICE1) expressed in the endosperm specifically downregulated JA synthesis and defense genes in the aleurone to promote starch synthesis, suggesting that regulatory units within the endosperm modulate defense vs growth through JA. We note that *HvAOC* is expressed in the aleurone at 7 DAP, in addition to maternal tissues, so could also control JA production in filial transfer tissues. Nonetheless, our evidence of spatial patterns of JA biosynthesis and signalling components coupled with segregation patterns indicate a central role for maternal tissue in generating JA and/or precursors or other products which themselves move to directly or otherwise drive signaling in both maternal and filial tissues to change grain dimensions (Fig 7). Altogether, we propose a mechanism, aligned with the classic growth versus defense paradigm (Guo *et al*., 2018), wherein JA limits maternal growth and accelerates its degeneration and nutrient delivery to regulate final grain parameters. As such, manipulating the JA pathway presents a possible route to increased grain size by lengthening the window through which parental resources can be transported to the filial tissues. Moreover, altering the rate of cell death or senescence during grain development could also help increase resiliency of grain fill under stressful conditions. Regulating jasmonate metabolism is an important mechanism used to promote wheat growth under elevated temperatures (Zhu *et al*., 2021). It will be important to learn whether similar mechanisms also work in the grain under warmer conditions.

## Acknowledgements

We are grateful for support from the University of Dundee Imaging Facility as well as Imaging Technologies at the James Hutton Institute (JHI). We also acknowledge support from the Barley Genetics group at JHI for providing germplasm and expertise, especially Pauline Smith, Richard Keith, Chris Warden, Dr Joanne Russell and Prof Robbie Waugh. We also acknowledge services and support from the Genome Technology Lab at JHI. This research was funded by Biotechnology and Biological Sciences Research Council grant BB/W003074/1 awarded to SMM. JL was supported by the China Scholarship Council and the University of Dundee. JG and JS were supported by the University of Dundee and BB/W003074/1. This research was facilitated by a UK-Australia Partnership Award BB/V018299/1 awarded to SMM. XY is supported by the Mortlock Bequest, Adelaide University. The authors acknowledge the instruments and expertise of Microscopy Australia (ROR: 042mm0k03) at Adelaide Microscopy, University of Adelaide, enabled by NCRIS, university and state government support.

## Author Contributions

SMM, JL and LL designed the research. JL, JS, MN, MKF, RH, EL and SMM performed the research. WW and JG developed experimental techniques. SMM, JL and LL analysed experimental data. SMM wrote the manuscript with contributions from JL. XY and MT shared VISIUM data and methods. All other authors had input into the manuscript.

## Data availability

All RNAseq data will be made available upon publication in publically available databases. Germplasm available upon request.

## Supporting Information

**Figure S1** Phylogenetic relationships and motif comparisons of AOC proteins from monocots and eudicots

**Figure S2** Allelic variation and haplotype structure of *HvAOC*.

**Figure S3** Expression patterns of *HvAOC* in different barley organs.

**Figure S4** Germination rate of Golden Promise (WT), *hvaoc-n4* and *hvaoc-n7* grain

**Figure S5** Spike-related phenotypes of Golden Promise (WT) *hvaoc-n4* and *hvaoc-n7*.

**Figure S6** Plant height of *Zeo1.b* x *hvaoc-n4* F3 siblings.

**Figure S7** Tissue-specific expression heatmaps of JA biosynthesis genes in developing barley grains.

**Figure S8** (a–b) Tissue-specific expression heatmaps of JA signalling components in developing barley grains.

**Table S1** Germplasm used in this study

**Table S2** Guide sequences for gene editing.

**Table S3** Primers used for genotyping.

**Table S4** Expected and observed segregation of *HvAOC* and *HvAP2* alleles in *Zeo1.b* x *hvaoc1-n4* F2 population and F3 siblings.

**Table S5** TUNEL positive spot counts of *hvaoc-n4* and Golden Promise at 6 and 12 days after pollination (DAP).

**Table S6** – Differentially expressed genes in *hvaoc-n4* compared to Golden Promise caryopses.

**Table S7** – Gene Ontogeny enrichment of differentially expressed genes

**Table S8** – heat map values of JA biosynthesis and signalling pathway

## References

1. Borisjuk L, Rolletschek H. 2009. The oxygen status of the developing seed. New Phytologist 182: 17–30.

2. Brinton J, Uauy C. 2019. A reductionist approach to dissecting grain weight and yield in wheat. Journal of Integrative Plant Biology 61: 337–358.

3. Campos ML, Kang J-H, Howe GA. 2014. Jasmonate-triggered plant immunity. Journal of Chemical Ecology 40: 657–675.

4. Chen C, Begcy K, Liu K, Folsom JJ, Wang Z, Zhang C, Walia H. 2016. Heat stress yields a unique MADS box transcription factor in determining seed size and thermal sensitivity. Plant Physiology 171: 606–622.

5. Domínguez F, Cejudo FJ. 2014. Programmed cell death (PCD): an essential process of cereal seed development and germination. Frontiers in Plant Science 5: 366.

6. Folsom JJ, Begcy K, Hao X, Wang D, Walia H. 2014. Rice fertilization-independent endosperm1 regulates seed size under heat stress by controlling early endosperm development. Plant Physiology 165: 238–248.

7. Guo Q, Major IT, Howe GA. 2018. Resolution of growth–defense conflict: mechanistic insights from jasmonate signaling. Current Opinion in Plant Biology 44: 72–81.

8. Hands P, Kourmpetli S, Sharples D, Harris RG, Drea S. 2012. Analysis of grain characters in temperate grasses reveals distinctive patterns of endosperm organization associated with grain shape. Journal of Experimental Botany 63: 6253–6266.

9. Hao J, Wang D, Wu Y, Huang K, Duan P, Li N, Xu R, Zeng D, Dong G, Zhang B, et al. 2021. The GW2-WG1-OsbZIP47 pathway controls grain size and weight in rice. Molecular Plant 14: 1266–1280.

10. Houston K, McKim SM, Comadran J, Bonar N, Druka I, Uzrek N, Cirillo E, Guzy-Wrobelska J, Collins NC, Halpin C. 2013. Variation in the interaction between alleles of HvAPETALA2 and microRNA172 determines the density of grains on the barley inflorescence. Proceedings of the National Academy of Sciences 110: 16675–16680.

11. Hu S, Yang H, Gao H, Yan J, Xie D. 2021. Control of seed size by jasmonate. Science China Life Sciences 64: 1215–1226.

12. Huang H, Liu B, Liu L, Song S. 2017. Jasmonate action in plant growth and development. Journal of Experimental Botany 68: 1349–1359.

13. Jia M, Li Y, Wang Z, Tao S, Sun G, Kong X, Wang K, Ye X, Liu S, Geng S, et al. 2021. TaIAA21 represses TaARF25-mediated expression of TaERFs required for grain size and weight development in wheat. The Plant Journal 108: 1754–1767.

14. Kellogg EA. 2016. Flowering Plants. Monocots. Springer, Switzerland: Springer Cham.

15. Kovacik M, Nowicka A, Zwyrtková J, Strejčková B, Vardanega I, Esteban E, Pasha A, Kaduchová K, Krautsova M, Červenková M, et al. 2024. The transcriptome landscape of developing barley seeds. The Plant Cell 36: 2512–2530.

16. Langer M, Hilo A, Guan J-C, Koch KE, Xiao H, Verboven P, Gündel A, Wagner S, Ortleb S, Radchuk V. 2023. Causes and consequences of endogenous hypoxia on growth and metabolism of developing maize kernels. Plant Physiology 192: 1268–1288.

17. Lawrenson T, Harwood WA 2018. Creating targeted gene knockouts in barley using CRISPR/Cas9. In: Harwood, W, eds. Barley: Methods and protocols. Springer, New York: Humana Press, 217–232.

18. Lawrenson T, Shorinola O, Stacey N, Li C, Østergaard L, Patron N, Uauy C, Harwood W. 2015. Induction of targeted, heritable mutations in barley and *Brassica oleracea* using RNA-guided Cas9 nuclease. Genome Biology 16: 258.

19. Lee SH, Sakuraba Y, Lee T, Kim KW, An G, Lee HY, Paek NC. 2015. Mutation of *Oryza sativa* CORONATINE INSENSITIVE 1b (OsCOI1b) delays leaf senescence. Journal of Integrative Plant Biology 57: 562–576.

20. Li M, Yu G, Cao C, Liu P. 2021. Metabolism, signaling, and transport of jasmonates. Plant Communications 2: e100231.

21. Li X, Wang Y, Duan E, Qi Q, Zhou K, Lin Q, Wang D, Wang Y, Long W, Zhao Z. 2018. OPEN GLUME1: a key enzyme reducing the precursor of JA, participates in carbohydrate transport of lodicules during anthesis in rice. Plant Cell Reports 37: 329–346.

22. Li Y, Wu F, Li C. 2025. Jasmonate signaling: integrating stress responses with developmental regulation in plants. Journal of Genetics and Genomics 52: 1490–1506.

23. Liu L, Jose SB, Campoli C, Bayer MM, Sánchez-Diaz MA, McAllister T, Zhou Y, Eskan M, Milne L, Schreiber M, et al. 2022. Conserved signalling components coordinate epidermal patterning and cuticle deposition in barley. Nature Communications 13: 6050.

24. Liu L, Zou Z, Qian K, Xia C, He Y, Zeng H, Zhou X, Riemann M, Yin C. 2017. Jasmonic acid deficiency leads to scattered floret opening time in cytoplasmic male sterile rice Zhenshan 97A. Journal of Experimental Botany 68: 4613–4625.

25. Liu X, Chi H, Yue M, Zhang X, Li W, Jia E. 2012. The regulation of exogenous jasmonic acid on UV-B stress tolerance in wheat. Journal of Plant Growth Regulation 31: 436–447.

26. Long Y, Wang C, Liu C, Li H, Pu A, Dong Z, Wei X, Wan X. 2024. Molecular mechanisms controlling grain size and weight and their biotechnological breeding applications in maize and other cereal crops. Journal of Advanced Research 62: 27–46.

27. Machado RA, McClure M, Herve MR, Baldwin IT, Erb M. 2016. Benefits of jasmonate-dependent defenses against vertebrate herbivores in nature. Elife 5: e13720.

28. Mehra P, Pandey BK, Verma L, Prusty A, Singh AP, Sharma S, Malik N, Bennett MJ, Parida SK, Giri J, et al. 2022. OsJAZ11 regulates spikelet and seed development in rice. Plant Direct 6: e401.

29. Milne L, Bayer M, Rapazote-Flores P, Mayer C-D, Waugh R, Simpson CG. 2021. EORNA, a barley gene and transcript abundance database. Scientific Data 8: 90.

30. Olsen O-A. 2004. Nuclear endosperm development in cereals and *Arabidopsis thaliana*. The Plant Cell 16: S214–S227.

31. Olsen O-A. 2020. The modular control of cereal endosperm development. Trends in Plant Science 25: 279–290.

32. Patil V, McDermott HI, McAllister T, Cummins M, Silva JC, Mollison E, Meikle R, Morris J, Hedley PE, Waugh R. 2019. APETALA2 control of barley internode elongation. Development 146: dev170373.

33. Peirats-Llobet M, Yi C, Liew LC, Berkowitz O, Narsai R, Lewsey MG, Whelan J. 2023. Spatially resolved transcriptomic analysis of the germinating barley grain. Nucleic Acids Research 51: 7798–7819.

34. Radchuk V, Belew ZM, Gündel A, Mayer S, Hilo A, Hensel G, Sharma R, Neumann K, Ortleb S, Wagner S. 2023. SWEET11b transports both sugar and cytokinin in developing barley grains. The Plant Cell 35: 2186–2207.

35. Radchuk V, Borisjuk L, Radchuk R, Steinbiss H-H, Rolletschek H, Broeders S, Wobus U. 2006. Jekyll encodes a novel protein involved in the sexual reproduction of barley. The Plant Cell 18: 1652–1666.

36. Radchuk V, Tran V, Hilo A, Muszynska A, Gündel A, Wagner S, Fuchs J, Hensel G, Ortleb S, Munz E. 2021. Grain filling in barley relies on developmentally controlled programmed cell death. Communications Biology 4(1): 428.

37. Radchuk V, Weier D, Radchuk R, Weschke W, Weber H. 2011. Development of maternal seed tissue in barley is mediated by regulated cell expansion and cell disintegration and coordinated with endosperm growth. Journal of Experimental Botany 62: 1217–1227.

38. Ren D, Hu J, Xu Q, Cui Y, Zhang Y, Zhou T, Rao Y, Xue D, Zeng D, Zhang G, et al. 2018. FZP determines grain size and sterile lemma fate in rice. Journal of Experimental Botany 69: 4853–4866.

39. Riemann M, Haga K, Shimizu T, Okada K, Ando S, Mochizuki S, Nishizawa Y, Yamanouchi U, Nick P, Yano M. 2013. Identification of rice Allene Oxide Cyclase mutants and the function of jasmonate for defence against *Magnaporthe oryzae*. The Plant Journal 74: 226–238.

40. Rolletschek H, Weschke W, Weber H, Wobus U, Borisjuk L. 2004. Energy state and its control on seed development: starch accumulation is associated with high ATP and steep oxygen gradients within barley grains. Journal of Experimental Botany 55: 1351–1359.

41. Shoesmith JR, Solomon CU, Yang X, Wilkinson LG, Sheldrick S, van Eijden E, Couwenberg S, Pugh LM, Eskan M, Stephens J. 2021. APETALA2 functions as a temporal factor together with BLADE-ON-PETIOLE2 and MADS29 to control flower and grain development in barley. Development 148: dev194894.

42. Tanabe S, Ashikari M, Fujioka S, Takatsuto S, Yoshida S, Yano M, Yoshimura A, Kitano H, Matsuoka M, Fujisawa Y, et al. 2005. A novel cytochrome P450 is implicated in brassinosteroid biosynthesis via the characterization of a rice dwarf mutant, dwarf11, with reduced seed length. The Plant Cell 17: 776–790.

43. Thiel J, Weier D, Sreenivasulu N, Strickert M, Weichert N, Melzer M, Czauderna T, Wobus U, Weber H, Weschke W. 2008. Different hormonal regulation of cellular differentiation and function in nucellar projection and endosperm transfer cells: A microdissection-based transcriptome study of young barley grains. Plant Physiology 148: 1436–1452.

44. Wang F, Lin J, Yang F, Chen X, Liu Y, Yan L, Chen J, Wang Z, Xie H, Zhang J, et al. 2024. The OsMAPK5–OsWRKY72 module negatively regulates grain length and grain weight in rice. Journal of Integrative Plant Biology 66: 2648–2663.

45. Wasternack C, Forner S, Strnad M, Hause B. 2013. Jasmonates in flower and seed development. Biochimie 95: 79–85.

46. Wasternack C, Hause B. 2013. Jasmonates: biosynthesis, perception, signal transduction and action in plant stress response, growth and development. An update to the 2007 review in Annals of Botany. Annals of Botany 111: 1021–1058.

47. Wasternack C, Kombrink E. 2010. Jasmonates: structural requirements for lipid-derived signals active in plant stress responses and development. ACS Chemical Biology 5: 63–77.

48. Wasternack C, Strnad M. 2018. Jasmonates: news on occurrence, biosynthesis, metabolism and action of an ancient group of signaling compounds. International Journal of Molecular Sciences 19: 2539.

49. Yang C-Y, Hsu F-C, Li J-P, Wang N-N, Shih M-C. 2011. The AP2/ERF transcription factor AtERF73/HRE1 modulates ethylene responses during hypoxia in Arabidopsis. Plant Physiology 156: 202–212.

50. Yang X, Wilkinson LG, Aubert MK, Houston K, Shirley NJ, Tucker MR. 2023. Ovule cell wall composition is a maternal determinant of grain size in barley. New Phytologist 237: 2136–2147.

51. Yin L-L, Xue H-W. 2012. The MADS29 transcription factor regulates the degradation of the nucellus and the nucellar projection during rice seed development. The Plant Cell 24: 1049–1065.

52. Zhao D-S, Li Q-F, Zhang C-Q, Zhang C, Yang Q-Q, Pan L-X, Ren X-Y, Lu J, Gu M-H, Liu Q-Q. 2018. GS9 acts as a transcriptional activator to regulate rice grain shape and appearance quality. Nature Communications 9: 1240.

53. Zhu T, Herrfurth C, Xin M, Savchenko T, Feussner I, Goossens A, De Smet I. 2021. Warm temperature triggers JOX and ST2A-mediated jasmonate catabolism to promote plant growth. Nature Communications 12: 4804.

