## Supplementary Information for "Control of grain dimensions via ALLENE OXIDE CYCLASE"

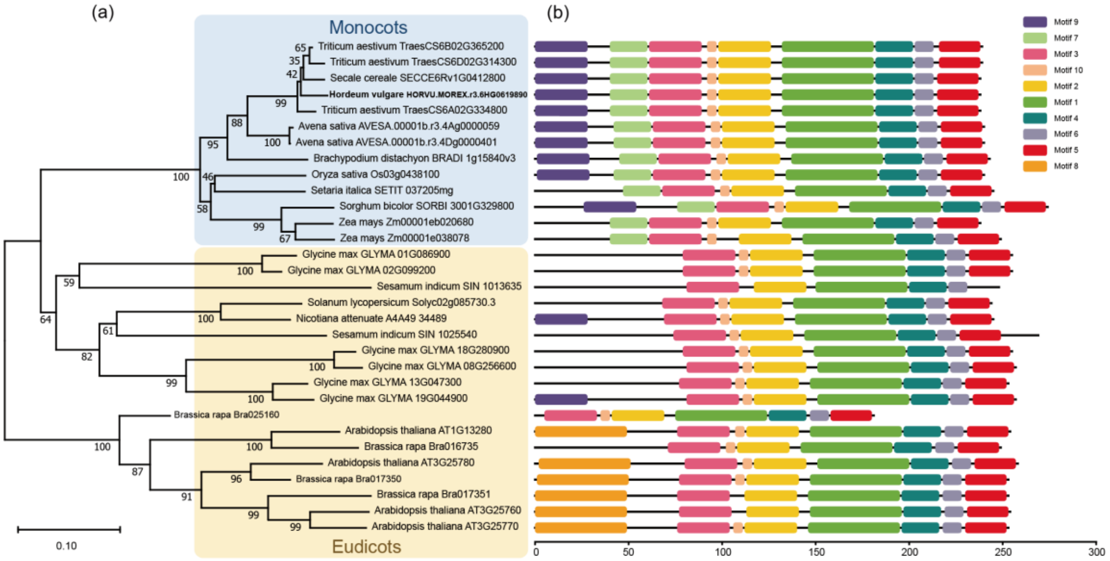


**Figure S1** Phylogenetic relationships and motif comparisons of AOC proteins from monocots and eudicots. (a) Maximum-likelihood phylogenetic tree of AOC protein sequences constructed from representative monocot (blue area) and eudicot (yellow area) species. Bootstrap values from 1000 replicates are shown at the nodes. (b) Conserved motif distribution of AOC proteins identified using MEME. Each colored box represents an individual motif.

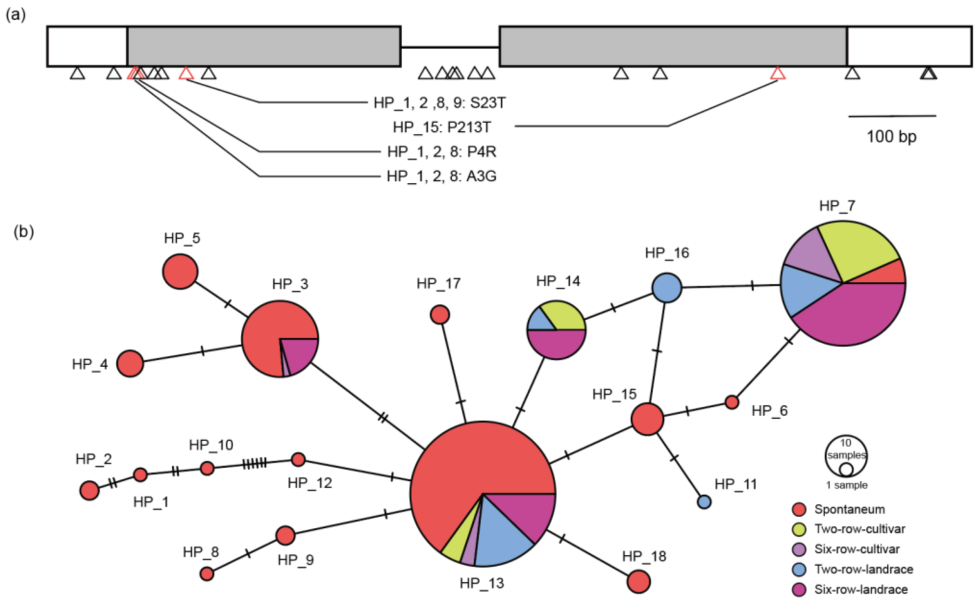


**Figure S2** Allelic variation and haplotype structure of *HvAOC*. (a) SNP sites within *HvAOC* identified from the whole-exome sequencing of 442 diverse barley accessions, including *Hordeum spontaneum* (wild barley), cultivars and landraces. Grey boxes represent exons and white boxes untranslated regions. Black triangles indicate synonymous or non-coding SNPs, while red triangles indicate non-synonymous SNPs sites. Haplotypes (HPs) associated with amino acid substitutions are indicated below the gene model. (b) Median-joining haplotype network of *HvAOC* in cultivated, landrace and wild barley. Each circle represents a haplotype, with circle size proportional to the haplotype frequency. Short bars on connecting lines indicate the number of mutational differences between HPs.

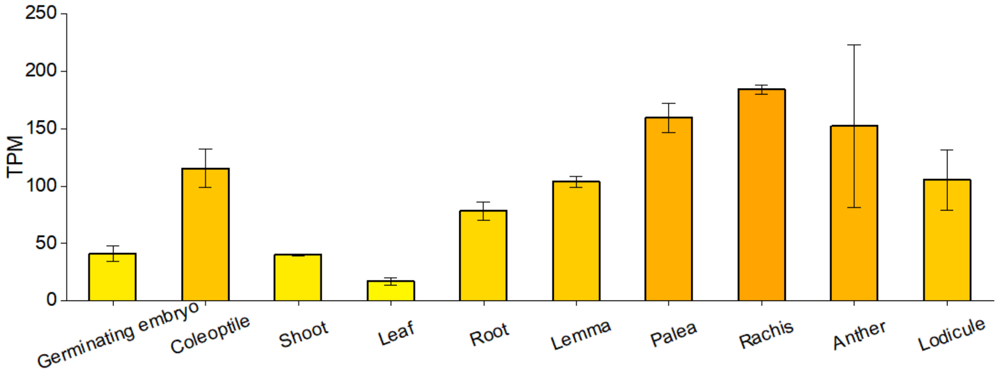


**Figure S3** Expression patterns of *HvAOC* in different barley organs. Level of *HvAOC* transcripts in various barley tissues. Bars on the top of the column represent standard deviation (n = 3). Expression data were obtained from EoRNA (https://ics.hutton.ac.uk/eorna/index.html). TPM, transcripts per million.

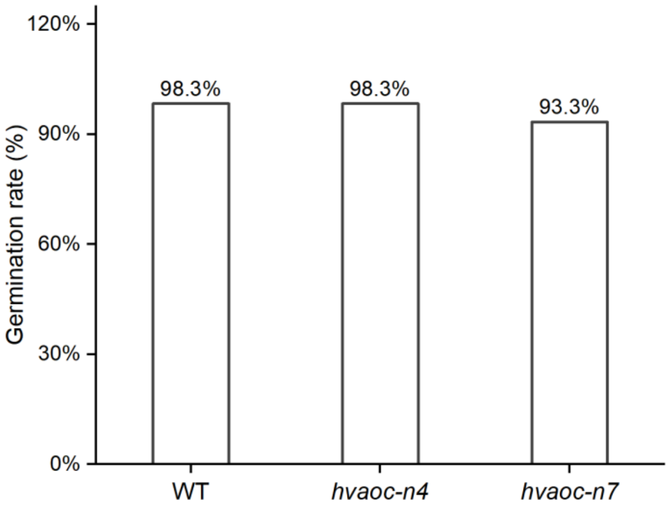


**Figure S4** Germination rate of Golden Promise (WT), *hvaoc-n4* and *hvaoc-n7* grain (n = 60/ genotype).

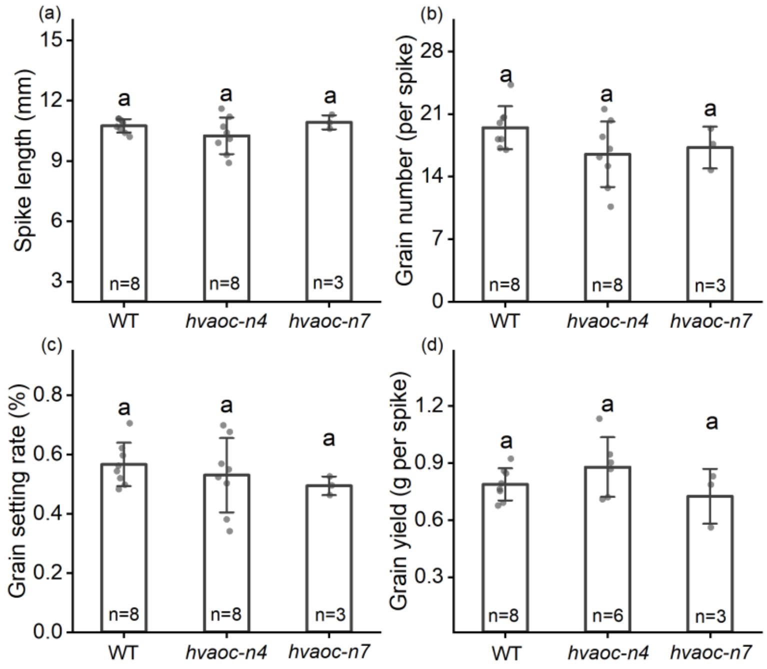


**Figure S5** Spike-related phenotypes of Golden Promise (WT) *hvaoc-n4* and *hvaoc-n7*. Data are shown as means ± SD (n ≥ 3), gray dots at the top of the bars represent individual biological replicates. Bars on the top of the column represent standard deviation and different letters indicate significant differences ($p < 0.05$, one-way ANOVA, Tukey's HSD test).

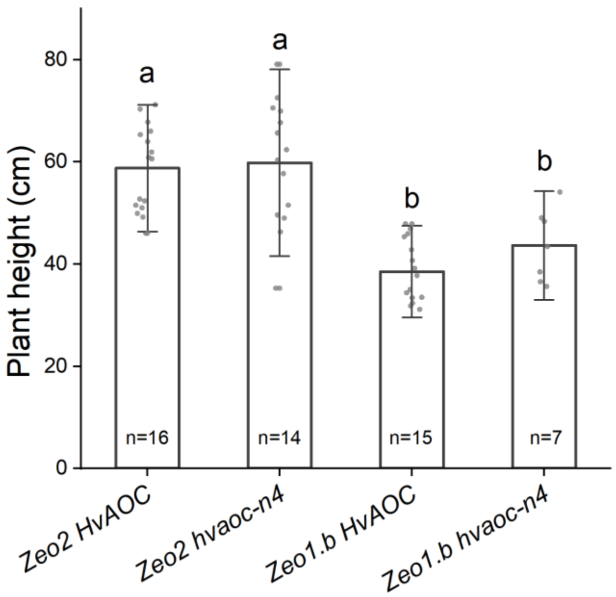


**Figure S6** Plant height of *Zeo1.b* x *hvaoc-n4* F3 siblings. Gray dots represent biological replicates (n≥7), w. Bars on the top of the column represent standard deviation and different letters indicate significant differences ($p < 0.05$, one-way ANOVA, Tukey's HSD test).

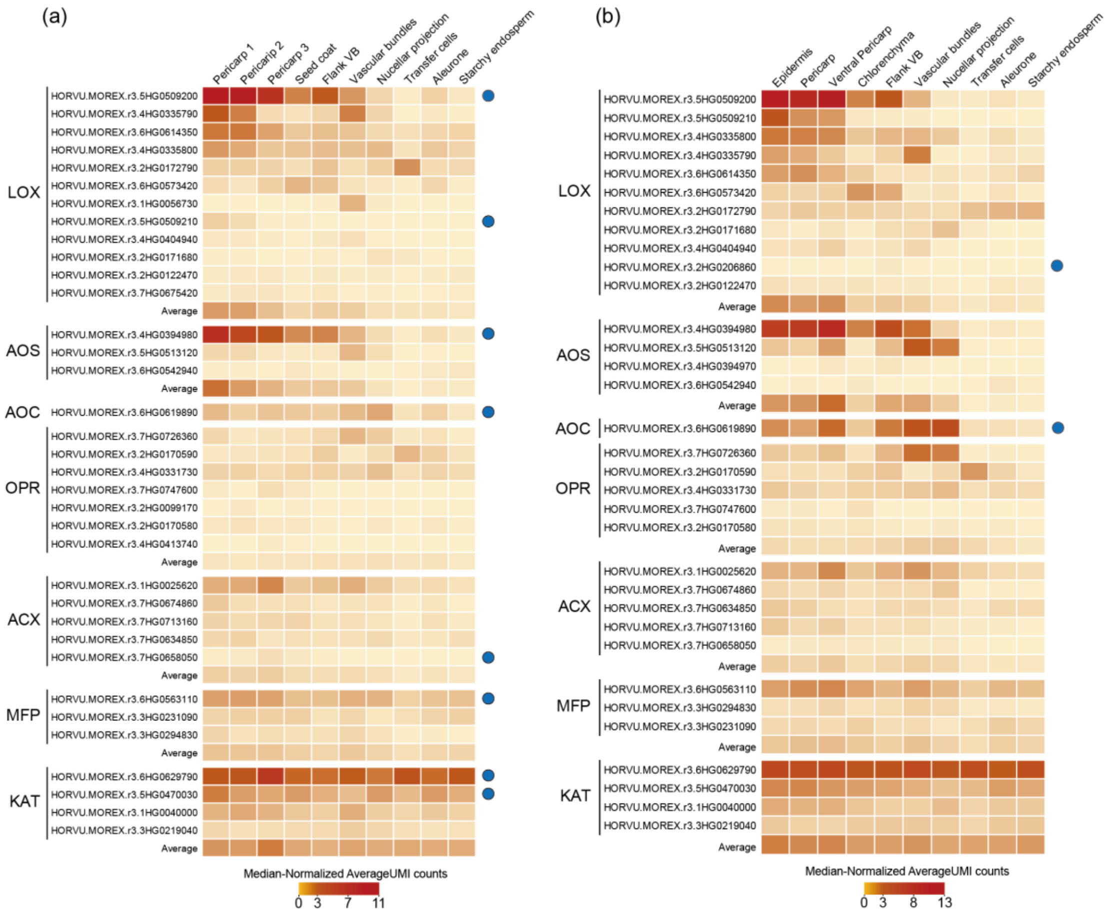


**Figure S7** Tissue-specific expression heatmaps of JA biosynthesis genes in developing barley grains. (a–b) JA biosynthetic genes expression across spatial transcriptomic tissue domains at 7 DAP (a) and 10 DAP (b). Blue circles in (a) and (b) indicate genes that were significantly downregulated (|$\log_2$ fold change|≥0.5 and a Benjamini–Hochberg adjusted p-value ≤ 0.05) in *hvaoc-n4* compared to WT in the bulk RNA-seq dataset at the 6 DAP (a) and 12 DAP (b), respectively. Color scale represents median-normalized average UMI counts. Genes with expression levels below 0.1 in all tissues were excluded from the heatmap. The complete raw data are provided in Supplementary Table S8.

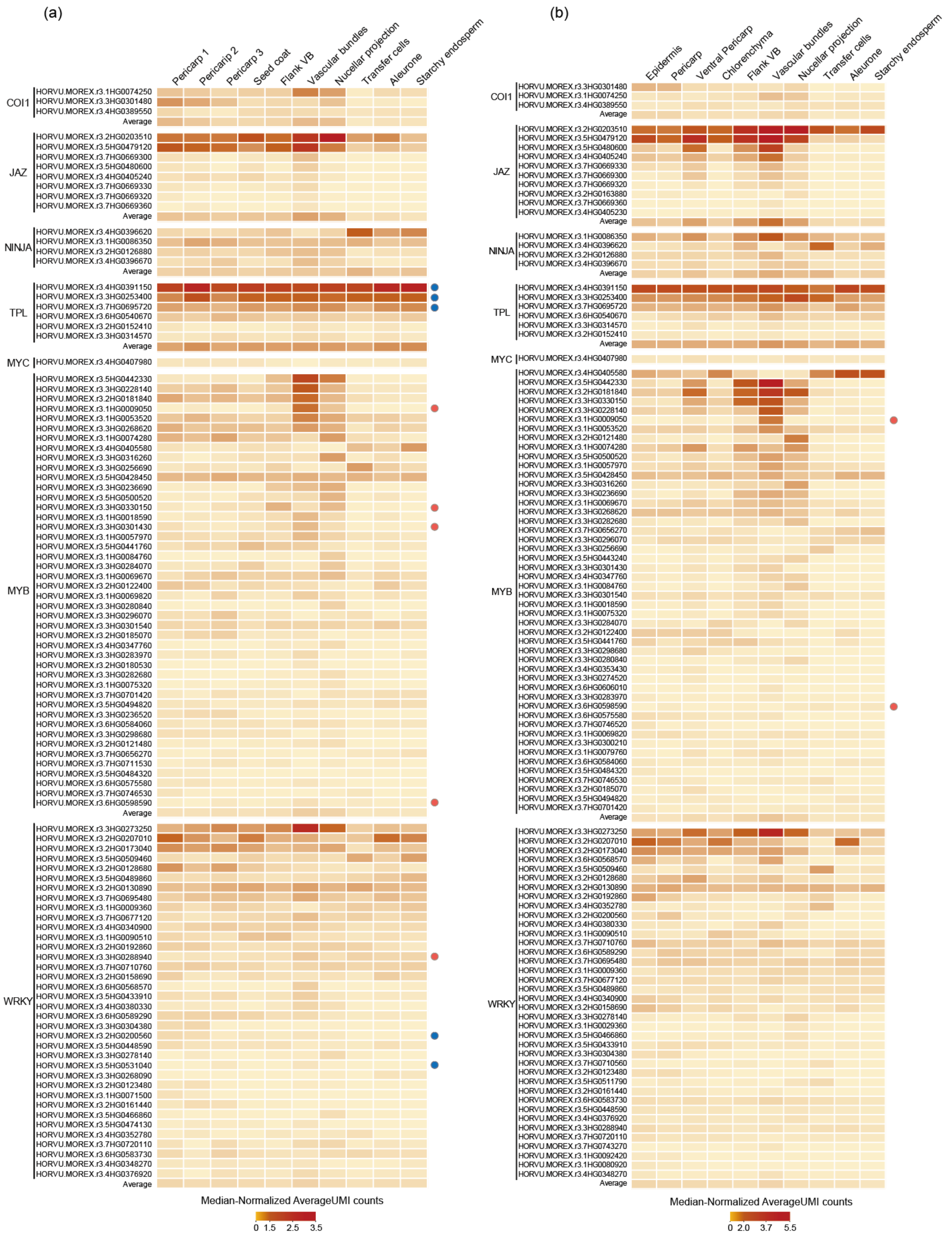

**Figure S8** Tissue-specific expression heatmaps of JA signalling components in developing barley grains. (a–b) JA signalling components gene expressions across spatial transcriptomic tissue domains at 7 DAP (a) and 10 DAP (b). Coloured circles in (a) and (b) indicate significant differential expressed genes (|$\log_2$ fold change|≥0.5 and a Benjamini–Hochberg adjusted p-value ≤ 0.05) in the *hvaoc-n4* relative to WT in bulk RNA-seq at 6 DAP and 12 DAP, respectively. Blue circles mean significantly downregulated gene, while red circles represent significantly upregulated genes in *hvaoc-n4* at the corresponding stage. Color scale represents median-normalized average UMI counts. Genes with expression levels below 0.1 in all tissues were excluded from the heatmap. The complete raw data are provided in Supplementary Table S8.

**Table S1** Germplasm used in this study.

| Allele | Mutagen | Cultivar | Molecular nature | Bowman Near Isogenic Line (backcross#) | Phenotype | Gene name HORVU gene model |
|---|---|---|---|---|---|---|
| *hvaoc-n4* | CRISPR-Cas9 | Golden Promise | 81 bp insertion & 17 bp deletion | n/a | Elongated coleoptile and protruding gran | HvAOC, HORVU.MOREX.r3.6HG0619890 |
| *hvaoc-n7* | CRISPR-Cas9 | Golden Promise | 1 bp insertion | n/a | Elongated coleoptile and protruding gran | HvAOC, HORVU.MOREX.r3.6HG0619890 |
| *Zeo1.b* | X-ray | Donaria | SNP a<g in mir172binding site | BWNIL938(BC5) | Dense spike, non-swelling lodicules | HvAP2, HORVU.MOREXr3.2HG0204770 |

**Table S2** Guide sequences for gene editing cloned into vector L2\Hv\35-36.

| | **Protospacer 5'-3'** | **sgDNA sequence 5'-3'** |
|---|---|---|
| HvAOC Guide 1a | gccgcgagggctgggctcgg | gttttagagctagaaatagcaagttaaaataaggctagtccgttatcaacttgaaaaagtg gcaccgagtcggtgcttttttttctagacccagctttcttgtacaaagttggcattacgctttac |
| HvAOC Guide 1b | gctcttctcgcccaagcctg | gttttagagctagaaatagcaagttaaaataaggctagtccgttatcaacttgaaaaagtg gcaccgagtcggtgcttttttttctagacccagctttcttgtacaaagttggcattacgctttac |

**Table S3** Primers used for genotyping.

| **Primer** | **Forward primer 5’-3’** | **Reverse Primer 5’-3’** |
|---|---|---|
| HvAOC | GCGTACCAGCTCATCGTAGT | TGAATCACTCAACGAAGCAGA |
| Cas9_1 | ACAAAGGCGGCAACAAACG | TAGCTGGGCAATGGAATCCG |
| Cas9_2 | TACGGACGAGTACAAGGTGC | GTCTGACAGGTTCTTTGCCG |
| Zeo1.b | AGCAACAGAGGCTGCAGCAC | GTTGGTTCGGCGGAACACAC |

**Table S5** Expected and observed segregation of *HvAOC* and *HvAP2* alleles in *Zeo1.b* x *hvaoc1-n4* F2 population and F3 siblings.

| Population | AOC allele | AP2 allele | Expected | Observed | Statistics |
|---|---|---|---|---|---|
| *Zeo1.b* x *hvaoc-n4* F2 population segregation | *hvaoc-n4/ hvaoc-n4* | *Zeo1.b / Zeo1.b* | 4 | 2 | χ2 =4.63; P = 0.80 |
| | *hvaoc-n4/ hvaoc-n4* | *Zeo1.b / Zeo2* | 8 | 7 | |
| | *hvaoc-n4/ HvAOC* | *Zeo1.b / Zeo1.b* | 8 | 6 | |
| | *hvaoc-n4/ HvAOC* | *Zeo1.b / Zeo2* | 15 | 18 | |
| | *hvaoc-n4/ hvaoc-n4* | *Zeo2 / Zeo2* | 4 | 3 | |
| | *hvaoc-n4/ HvAOC* | *Zeo2 / Zeo2* | 12 | 12 | |
| | *HvAOC / HvAOC* | *Zeo2 / Zeo2* | 4 | 4 | |
| | *HvAOC / HvAOC* | *Zeo1.b / Zeo1.b* | 4 | 5 | |
| | *HvAOC / HvAOC* | *Zeo1.b / Zeo2* | 4 | 7 | |
| *Zeo1.b/ Zeo1.b hvaoc-n4/ HvAOC* F3 family segregation | *hvaoc-n4/ hvaoc-n4* | *Zeo2 / Zeo2* | 15.75 | 17 | χ2 = 0.40; P = 0.82 |
| | *hvaoc-n4/ HvAOC* | *Zeo2 / Zeo2* | 31.5 | 29 | |
| | *HvAOC / HvAOC* | *Zeo2 / Zeo2* | 15.75 | 17 | |
| *Zeo2 / Zeo2 hvaoc-n4/ HvAOC* F3 family segregation | *hvaoc-n4/ hvaoc-n4* | *Zeo1.b / Zeo1.b* | 15 | 13 | χ2 = 0.37; P = 0.83 |
| | *hvaoc-n4/ HvAOC* | *Zeo1.b / Zeo1.b* | 30 | 31 | |
| | *HvAOC / HvAOC* | *Zeo1.b / Zeo1.b* | 15 | 16 | |

**Table S5** TUNEL positive spot counts of *hvaoc-n4* and Golden Promise at 6 and 12 days after pollination (DAP).

| **Days after pollination (DAP)** | **Tissue area** | **TUNEL positive spots counts** | |
|---|---|---|---|
| | | ***hvaoc-n4*** | **Golden Promise** |
| 6 DAP | FVB adjacent area | 12.3±3.4 | 35±5.0 |
| | VB | 10.2±11.4 | 21.0±5.4 |
| | NP | 29.3±10.5 | 35.4±3.6 |
| 12 DAP | FVB adjacent area | 14.1±1.9 | 30.1±6.9 |
| | VB | 10.9±2.5 | 27.2±10.4 |
| | NP | 32.2±18.8 | 49.9±18.0 |

Note: Data are presented as mean ± standard deviation (SD). Values were obtained from three biological replicates for both the *hvaoc-n4* mutant and wild-type (Golden Promise) and each replicates contained three individual grain sections as technical replicate. The FVB-adjacent area represents the sum of the positive TUNEL spots in left and right flanking vascular bundle regions. FVB, flanking vascular bundle; VB, vascular bundle; NP, nucellar projection.
